# Inflammation increases the penetrance of behavioral impairment in *Shank3* haploinsufficiency mice – can it explain the behavioral regression in Autism?

**DOI:** 10.1101/2025.08.07.669241

**Authors:** Sheng-Nan Qiao, Sung Eun Wang, Kun-Yong Kim, Sungsin Jo, Yong-Hui Jiang

## Abstract

Behavioral regression occurs in ∼40% of *SHANK3*-associated autism spectrum disorder (ASD). We previously reported that significant behavioral regression in a small cohort with *SHANK3* haploinsufficiency patients, triggered by subclinical infections, responded to immunomodulator treatments. We hypothesize that behavioral regression results from the interplay between *SHANK3* deficiency and neuroinflammation. Using *Shank3* exon 4-22 deletion heterozygous mutant (*Sh3^+/-^*) mouse, which shows no significant behavior impairments, we established a preclinical model – *Shank3* haploinsufficiency mouse undergoing systemic inflammation challenge via intraperitoneal injection of lipopolysaccharides (LPS). We found that, two weeks after LPS challenge, wild-type mice (WT) recovered but *Sh3^+/-^* mice exhibited motor impairment, anxiety-like behaviors, and excessive grooming, similar to *Shank3* exon 4-22 deletion homozygous mutants. Anti-inflammatory treatment partially reversed LPS-induced behavioral changes. Transcriptomic analysis revealed upregulation of neuroinflammation-related genes and downregulation of synaptic function-related genes in *Sh3^+/-^* mice in response to LPS. Especially, pro-inflammatory genes and microglia markers were overly activated that may result from the increased toll-like receptor 4 (TLR4) in *Sh3^+/-^* mice. Microglia overactivation elevated synapse engulfment and disrupted synaptic protein may underlie LPS-triggered worsen behavior phenotypes in *Sh3^+/-^* mice. Our findings indicate that neuroinflammation increases the penetrance of behavioral impairment in *Shank3* haploinsufficiency mice and support a potential mechanism for the behavioral regression in human *SHANK3* disorders for future investigations.

## Introduction

Autism spectrum disorder (ASD) is a group of neurodevelopmental conditions with typical onset before the age of three ^1^. A significant proportion of children with ASD shows developmental arrest at an early developmental age and behavioral regression throughout the lifespan. Behavioral regression is characterized by loss of previously acquired skills, including loss of purposeful hand use, loss of acquired speech, and gait abnormalities ^2–4^. Catatonic features are frequently associated with behavioral regression observed in ASD ^5,6^. However, the underlying mechanism for the early developmental arrest and behavioral regression remains poorly understood.

The *SHANK3* mutation is one of the most common monogenic defects associated with ASD in humans ^7–10^. Genetic mutations in *SHANK3* gene can manifest as Phelan-McDermid Syndrome (PMS), characterized by a chromosomal deletion of 22q13.3 or single nucleotide variants, indels, exonic deletions within the *SHANK3* gene in individuals with primary ASD ^11^. Patients with *SHANK3* haploinsufficiency also exhibit severe intellectual disability, motor deficits, and speech delays ^12–14^. Developmental regression with catatonic features has been reported in ∼40% of cases with *SHANK3* haploinsufficiency ^15–20^. We previously described four patients with pathogenic *SHANK3* mutations who experienced acute and severe behavioral regression following possible subclinical viral infections ^21^. Notably, treatment with intravenous immunoglobulins (IVIG) combined with anti-inflammatory drugs and immunomodulators significantly reversed the behavioral regression in these cases ^21^. This clinical report strongly suggests a potential link between inflammation and behavioral regression associated with *SHANK3* haploinsufficiency. However, the direct causality could not be investigated and determined from human studies.

SHANK3 protein is primarily characterized as a scaffold protein located at the postsynaptic density of neurons, interacting with other synaptic proteins and affecting synaptic function ^22^. However, the role of SHANK3 in function of non-neuronal cells such as microglia has been poorly studied ^23–26^. Microglia activation can sculpt synapses and brain circuits and change animal behavioral phenotypes during normal development as well as in response to neuroinflammation ^27,28^. Aberrant microglia activation has been associated with ASD^29–32^. Our previous studies showed that inflammation induces acute behavioral changes ^33^ and aggravates hypothermia in *Shank3* mutant mouse model ^34^. These initial findings suggest a potential mechanistic link between *SHANK3* deficiency and inflammatory response.

Our lab previously generated a *Shank3* mutant mouse line with a deletion of exon 4 to 22 (*Shank3^Δe^*^4–22^) that disrupts most of the *Shank3* coding region^35^. *Shank3^Δe^*^4–22^ homozygous but not heterozygous mice showed robust face validity for autism-like behaviors^35^. Therefore, *Shank3*^Δe4–22^ heterozygous mice offer a unique opportunity to examine the interaction between genes and environment, specifically whether neuroinflammation may modify the penetrance of behavioral phenotypes associated with *SHANK3* deficiency in humans. In this study, we utilized the *Shank3*^Δe4–22^ heterozygous mice and investigated the behavioral outcome, transcriptomic profiles, microglia activation, and synapse marker after systemic inflammatory challenge induced by lipopolysaccharides (LPS). These findings provide a general insight on the role of neuroinflammation in ASD susceptibility.

## Results

### Systemic inflammation via LPS modified the penetrance of motor dysfunction, anxiety-like behavior, and repetitive behavior in *Shank3*^Δe^^4–22^ heterozygous (*Sh3^+/-^*) mice

To explore whether inflammation plays a role in regression of ASD-associated behaviors, we challenged *Sh3^+/-^* mice by intraperitoneal injection of LPS (*Sh3^+/-^*+ LPS). Control mice were given the same volume of normal saline (PBS) (*Sh3*^+/-^ + PBS). Both LPS-treated *Sh3^+/-^*and wild-type (WT) mice experienced weight loss at 24 hours compared to PBS-treated mice (**Fig. 1A**). This indicated systemic inflammation induced by LPS, consistent with previous reports ^36^. LPS-treated mice regained body weight at 72 hours post-injection, suggesting recovery from the LPS challenge (**Fig. 1A**). Acute behavioral changes, including decreased locomotor activity ^37^, social avoidance ^38^, and depressive-like behavior ^39^, have been reported in LPS-treated mice around 24 hours post-injection. However, interpretating these behavioral changes in the context of ASD model may be confounded by immediate sickness of test mice ^33,40,41^. Therefore, we performed behavioral analysis two weeks after LPS injection to distinguish “LPS and SHANK3 haploinsufficiency interaction effect” vs. “acute LPS effect” (**Fig. 1B**). The *Shank3*^Δe4–22^ homozygous (*Sh3^-/-^*) group was included for a comparison ^35^.

**Figure 1.**
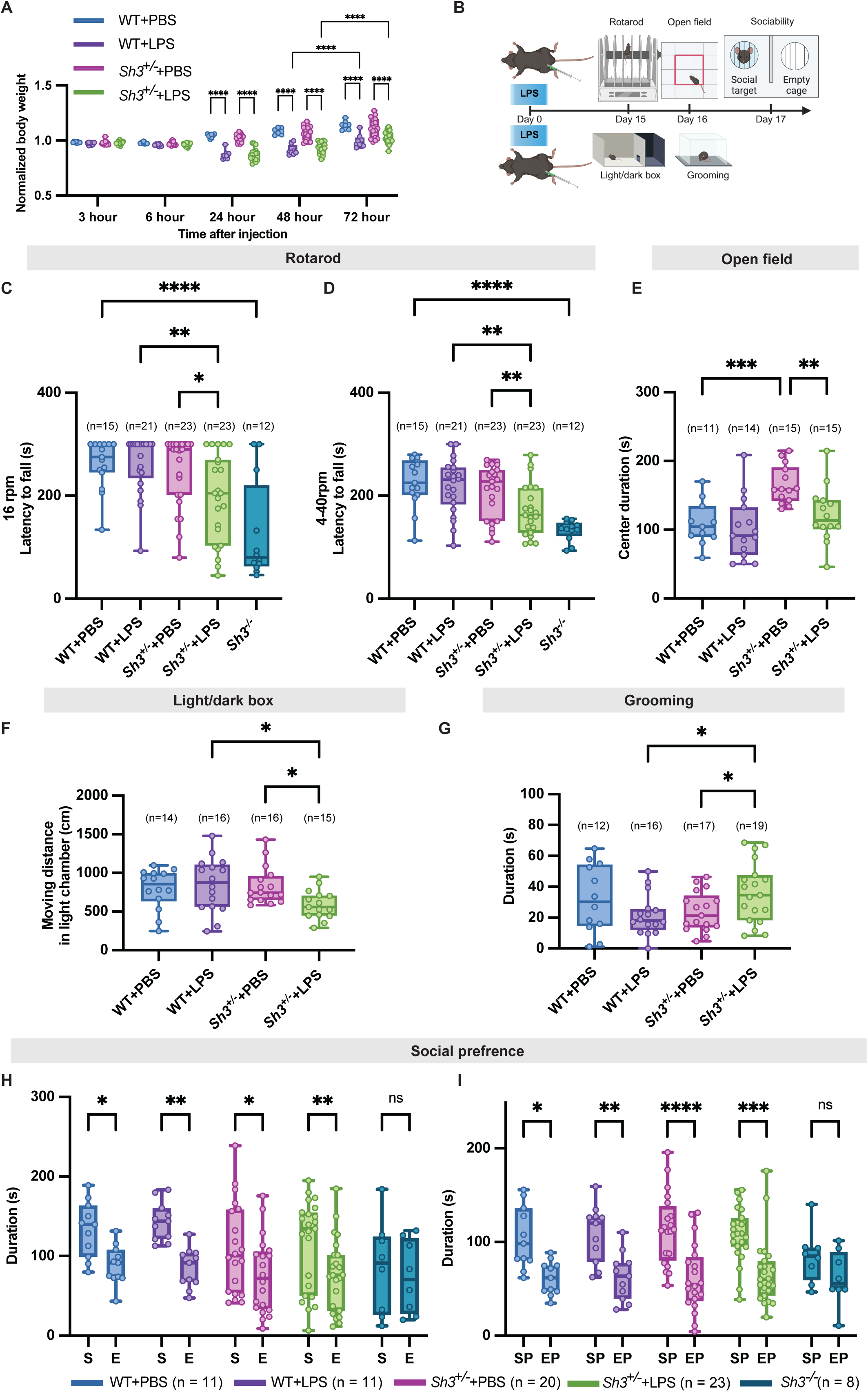
LPS challenge induced selective behavioral impairments in *Sh3^+/-^* mice. **A. The body weight loss after LPS intraperitoneal injection**. Mouse was weighed before injection, and at 3, 6, 24, 48, and 72 hours after LPS (1 mg/kg) or the same volume of PBS injection. Each mouse’s body weight was normalized to the weight before injection. Two-way repeated measures ANOVA was performed (genotype/LPS effect F_(3, 50)_ = 34.32, *p* < 0.0001; Hour effect F _(4, 200)_ = 84.15, *p* < 0.0001; genotype/LPS X Hour interaction F_(12, 200)_ = 22.64, *p* < 0.0001). Post-hoc Tuckey’s test showed no significant difference between WT+PBS (n = 8) and *Sh3^+/^*^-^+PBS (n = 17) or between WT+LPS (n = 8) and *Sh3^+/-^*+LPS (n = 21) at 3 hours, 6 hours, 24 hours, 48 hours and 72 hours after injection. Mouse body weight started to reduce significantly after LPS injection compared to PBS injection at 24 hours, 48 hours and 72 hours. Mouse body weight from LPS groups significantly increased at 72 hours compared to 48 hours after injection in both WT+LPS and *Sh3^+/^*^-^+LPS groups. See multiple comparison statistics in Supplementary Table 5. **B. Schematic drawing for the experimental design**. C-D. LPS challenge impaired motor function in *Sh3^+/-^* mice. Steady speed rotarod (16 rpm, **C**) and accelerated speed rotarod (4-40 rpm, **D**) tests were performed and latency to fall was measured. No significant difference was found between WT+PBS and *Sh3^+/^*^-^+PBS (16 rpm, *p* =0.5487, 4-40 rpm, *p* =0.3283) or between WT+PBS and WT+LPS (16 rpm, *p* =0.9797, 4-40 rpm, *p* =0.6711). *Sh3^+/-^*+LPS significantly reduced latency to fall compared to *Sh3^+/-^*+PBS (16 rpm, *p* =0.0126, 4-40 rpm, *p* =0.0061) or compared to WT+LPS (16 rpm, *p* =0.002, 4-40 rpm, *p* =0.001). *Sh3^-/-^* mice significantly reduced latency to fall compared to WT+PBS (16 rpm, *p* < 0.0001, 4-40 rpm, *p* < 0.0001). **E-F. LPS challenge increased anxiety-like behavior in *Sh3^+/-^* mice.** Mouse duration spent in the center of the open field test and travel distance in light chamber of light/dark box test were measured. No significant difference found between WT+PBS and WT+LPS (OFT, *p* = 0.5956, light/dark box *p* = 0.6004). *Sh3^+/-^+*LPS showed less center duration than *Sh3^+/-^+*PBS (*p* = 0.0033) despite that *Sh3^+/-^+*PBS showed increased center duration than WT+PBS (*p* = 0.0005), and *Sh3^+/-^+*LPS showed less travel distance in light chamber than *Sh3^+/-^+*PBS (*p* = 0.0158) or WT+LPS (*p* = 0.0107). **G. LPS challenge increased repetitive behavior in *Sh3^+/-^* mice.** Grooming behavior was hand scored and compared. *Sh3^+/-^+*LPS showed longer grooming duration than *Sh3^+/-^+*PBS (*p* = 0.0348) or WT+LPS (*p* = 0.0135). **H-I. LPS challenge had no effect on sociability *Sh3^+/-^* mice.** Three-chamber tests were conducted and duration spent in mouse chamber (S), empty chamber (E), proximity regions in mouse chamber (SP) and empty chamber (EP) were compared (Two-way repeated measures ANOVA. **H.** genotype effect F_(4, 68)_ = 1.590, *p* = 0.1869, S/E effect F _(1, 68)_ = 25.46, *p* < 0.0001, genotype X S/E interaction F_(4, 68)_ = 0.8135, *p* = 0.5209; **I.** genotype effect F _(4, 68)_ = 1.398, *p* = 0.2441, SP/EP effect F_(1, 68)_ = 34, *p* < 0.0001, genotype X SP/EP interaction F_(4, 68)_ = 0.4236, *p* = 0.7911). No significant difference of duration was found between S vs. E (*p* = 0.5325) or SP vs. EP (*p* = 0.2529) in *Sh3^-/-^* mice. All other groups of mice showed significantly increased duration in S vs. E or SP vs. EP. * *p* < 0.05, _**_ *p* < 0.005, *** *p* < 0.0005, **** *p* < 0.0001

To assess the motor coordination, latency to fall from the rotarod was measured in both steady speed (16 rpm) and accelerated speed (4-40 rpm) conditions. As expected, *Sh3^-/-^* mice showed significantly reduced latency to fall compared to WT+PBS group. No significant difference was observed between WT+PBS and WT+LPS groups, but *Sh3^+/-^*mice treated with LPS showed significantly shorter latency to fall than the PBS group (**Fig. 1C-D**). The duration of mouse spent in center of open field test (OFT) showed no significant difference between WT+PBS and WT+LPS groups, but *Sh3^+/-^*+LPS spent significantly less time in the center compared to *Sh3^+/-^*+PBS (**Fig. 1E**). In the light/dark box test, distance of mouse moved in light chamber showed no significant difference between WT+PBS and WT+LPS groups while *Sh3^+/-^*+LPS significantly moved less in light chamber compared to *Sh3^+/-^*+PBS (**Fig. 1F**). Repetitive behavior was assayed through scoring grooming behavior. *Sh3^+/-^*+LPS showed significantly longer grooming duration than *Sh3^+/-^*+PBS while WT+PBS and WT+LPS groups were comparable (**Fig. 1G**). We also conducted a three-chamber social preference test. As expected, *Sh3^-/-^* mice showed no preference for social targets compared to WT. However, both LPS- and PBS-treated *Sh3^+/-^* and WT mice spent significantly more time in the chamber with the social target (S) or in the proximity region of the social target (SP) (**Fig. 1H-I**).

To explore potential sex effect, we compared motor function and sociability in both PBS- and LPS-treated female and male *Sh3^+/-^* mice. LPS-treated male *Sh3^+/-^*mice exhibited significant reduced latency to fall in rotarod, although no significant difference found between LPS and PBS group in female *Sh3^+/-^* mice (**Supplementary Fig. 1A-B**). Both LPS- and PBS-treated male *Sh3^+/-^* mice spent significantly more time in S or SP, but no significant difference found in female *Sh3^+/-^*mice (**Supplementary Fig. 1C-D**). Taken together, our results indicated that, LPS treatment in *Sh3^+/-^* mice significantly impaired the motor performance on rotarod, changed exploratory or anxiety-like behaviors in OFT and light/dark box test, and increased grooming duration, but did not affect sociability in three-chamber test. By comparison, LPS treatment in WT mice did not have any significant effect on those behaviors. These findings suggest the interplay of inflammation and *Shank3* haploinsufficiency in select domains of behaviors.

To assess whether anti-inflammation treatment can reverse the behavioral changes induced by LPS in *Sh3^+/-^* mice, we applied mefenamic acid (MFA), a nonsteroidal anti-inflammatory drug (NSAID). One of the MFA targets is the intracellular cyclooxygenase (COX) signaling, which can be activated by LPS ^42–44^. Previous study also reported that MFA has neuroprotective effect ^45^. Given that both WT+LPS and *Sh3^+/-^* +LPS group exhibited impaired motor function at 1 week after the LPS injection (**Supplementary Fig. 2A-B**) but such behavior deficits of WT+LPS group recovered to baseline at 2-week post-injection (**Fig. 1C-D**), we chose day 7 post-injection as the endpoint of MFA treatment (**Fig. 2A**). We administered MFA via intraperitoneal injection once a day for seven days following LPS injection and then performed rotarod test 24 hours after the final dose of MFA to evaluate the treatment efficacy (**Fig. 2A**). In the steady speed (16 rpm) rotarod test, both WT+LPS+MFA and *Sh3^+/-^*+LPS+MFA groups showed significantly increased latency to fall compared to vehicle-treated groups (**Fig. 2B**), suggesting that MFA effectively mitigated the motor function deficits induced by LPS. *Sh3^+/-^*+LPS+MFA showed trend of shorter latency to fall than WT+LPS+MFA (**Fig. 2B**, *p* = 0.091), suggesting potential less pronounced anti-inflammatory efficacy and the presence of exacerbated inflammatory responses in *Sh3^+/-^* mice. In contrast, in the accelerated speed (4-40 rpm) rotarod test, MFA did not restore motor function in either WT or *Sh3^+/-^* mice (**Fig. 2C**). Moreover, when comparing MFA treatment in female and male *Sh3^+/-^* mice, MFA significantly increased latency to fall in female *Sh3^+/-^* mice but not in male *Sh3^+/-^* mice (**Fig. 2D-E**). Overall, our results showed that LPS-induced motor function impairment in *Sh3^+/-^* mice can be partially reversed by anti-inflammatory treatment. Future studies will include additional behavioral assays, as well as time-course and dose-dependent responses to MFA treatment.

**Figure 2.**
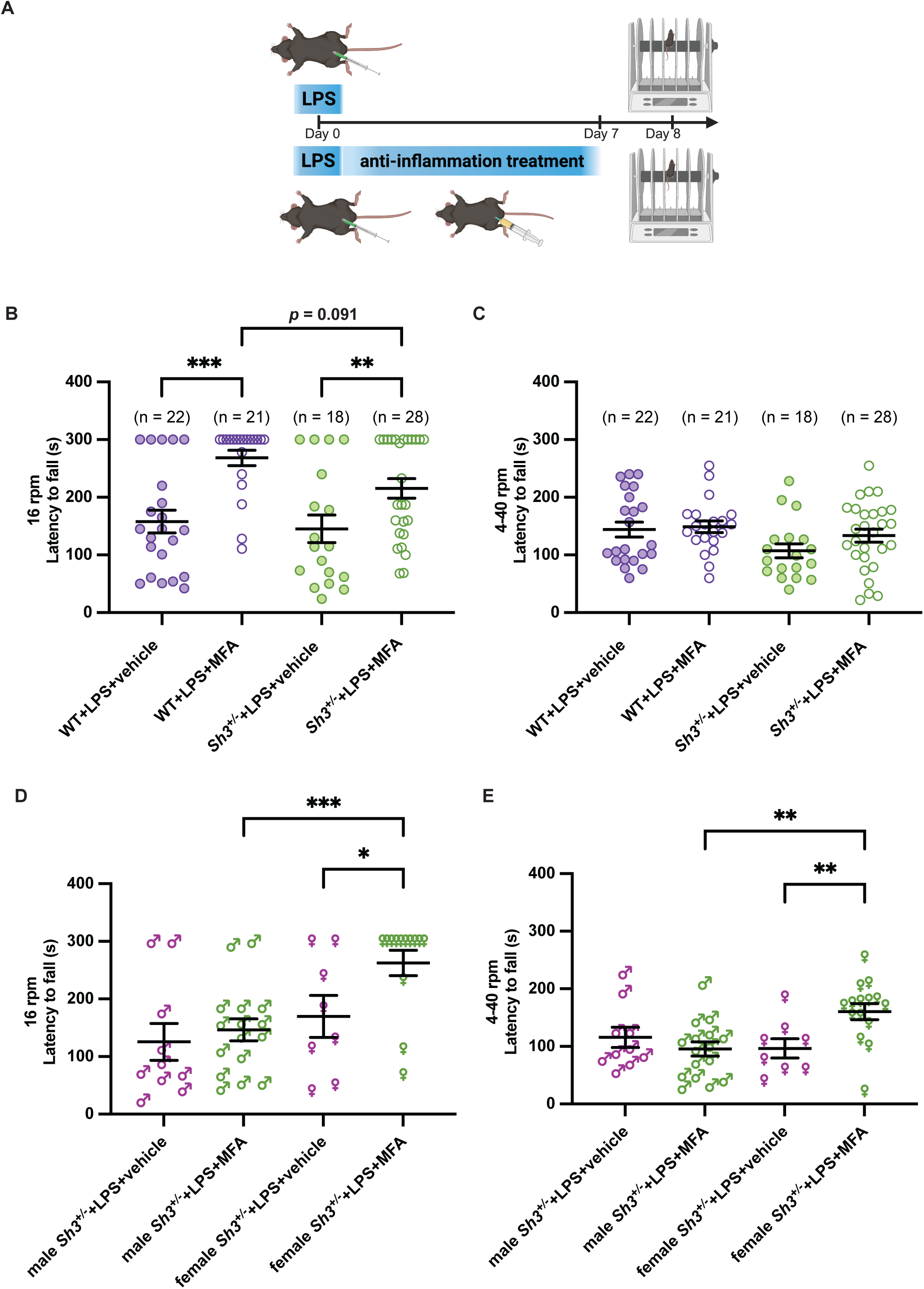
Mefenamic acid (MFA) partially improved the motor behavior deterioration in WT and *Sh3^+/-^* mice. **A. Schematic drawing for experiment design**. Anti-inflammatory treatment with intraperitoneal MFA injection was given to mice once every day for 7 days after LPS injection. Rotarod tests were performed 24 hours after the last does of MFA injection. **B-C. MFA partially improved the rotarod test performance.** After 1-week treatment of MFA, Steady speed rotarod (16 rpm, **B**) and accelerated speed rotarod (4-40 rpm, **C**) tests were performed and latency to fall was compared. (Two-way ANOVA, **B. 16 rpm,** LPS/MFA effect F _(1, 37)_ = 23.66, *p* < 0.0001, genotype effect F _(1,46)_ = 1.859, *p* = 0.1794, LPS/MFA X genotype interaction F _(1, 37)_ = 0.8697, *p* = 0.3571. **C. 4-40 rpm,** LPS/MFA effect F _(1, 85)_ = 1.721, *p* = 0.1932, genotype effect F _(1,85)_ = 4.822, *p* = 0.0308, LPS/MFA X genotype interaction F _(1, 85)_ = 0.8220, *p* = 0.3672). Compared to vehicle group, MFA-treated *Sh3^+/^*^-^ and WT mice showed increased latency to fall in 16 rpm rotarod test (*Sh3^+/^*^-^, *p* = 0.0091; WT, *p* = 0.0002). **D-E. MFA treatment was more effective in female *Sh3^+/-^* mice than male ones.** Two-way ANOVA was performed to compare sex difference in MFA treatment (**D. 16 rpm,** LPS/MFA effect F _(1, 43)_ = 4.626, *p* = 0.0372, sex effect F _(1,43)_ = 9.227, *p* = 0.004, LPS/MFA X sex interaction F _(1, 43)_ = 1.858, *p* = 0.1799. **E. 4-40 rpm,** LPS/MFA effect F _(1, 16)_ = 2.065, *p* = 0.17, sex effect F _(1,29)_ = 2.369, *p* = 0.1346, LPS/MFA X sex interaction F _(1, 16)_ = 8.046, *p* = 0.0119). MFA-treated female *Sh3^+/^*^-^ showed significantly higher latency to fall compared to vehicle-treated female *Sh3^+/^*^-^ (16 rpm, *p* = 0.0229; 40rpm, *p* = 0.0106) and MFA-treated male *Sh3^+/^*^-^ (16 rpm, *p* = 0.0009; 40rpm, *p* = 0.0011). * *p* < 0.05, ** *p* < 0.005, _***_ *p* < 0.0005, **** *p* < 0.0001

### LPS-induced differentially expressed genes (DEGs) were associated with neuroinflammation and neurotransmission in *Sh3^+/-^* mice

To understand how LPS-induced inflammation modifies the penetrance of motor impairment in *Sh3^+/-^* mice, we analyzed the transcriptomic profiles in the forebrains by bulk RNA-sequencing (RNA-seq) (**Supplementary Fig. 3A-B, Supplementary Tables 1 and 2**). LPS treatment induced hundreds of differentially expressed genes (DEGs) in both WT and *Sh3^+/-^* mice compared to PBS-treated groups. The number of upregulated genes (488 DEGs in *Sh3^+/-^*+LPS vs. *Sh3^+/-^*+PBS and 433 DEGs in WT+LPS vs. WT+PBS) was greater than that of downregulated genes (75 DEGs in *Sh3^+/-^*+LPS vs. *Sh3^+/-^*+PBS and 108 DEGs in WT+LPS vs. WT+PBS) (**Fig. 3A-B**). We identified 88 upregulated and 37 downregulated DEGs between *Sh3^+/-^*+LPS vs. WT+LPS (**Fig. 3C**). Although we found 67 upregulated and 33 downregulated genes when comparing transcriptomic profiles between *Sh3^+/-^*+PBS and WT+PBS (**Fig. 3D**), those genes barely overlapped with any LPS-induced DEGs observed in *Sh3^+/-^*mice. Next, we compared the overlapping genes among four comparisons (**Fig. 3E-F, Supplementary Tables 3 and 4**). A total of 313 shared upregulated and 12 shared downregulated DEGs were identified between WT+LPS vs. WT+PBS and *Sh3^+/-^*+LPS vs. *Sh3^+/-^*+PBS. Those shared DEGs probably represent the LPS-induced neuroinflammatory responses in mouse brains. Eight upregulated genes shared among *Sh3^+/-^*+LPS vs. WT+LPS, *Sh3^+/-^*+LPS vs. *Sh3^+/-^*+PBS and WT+LPS vs. WT+PBS were well-known for their roles in neuroinflammation (**Fig. 3E** highlighted with white underline, **Table 1**). Interestingly, 10 upregulated genes and 5 downregulated genes were found only in *Sh3^+/-^* mice not in WT mice in response to LPS (**Fig. 3E-F** highlighted with green underline, **Table 2-3**), suggesting that LPS-induced neuroinflammation in *Sh3^+/-^* mice is mediated by distinct signaling pathways. Additionally, when we compared the overlapping genes between *Sh3^+/-^*+PBS vs. WT+PBS and *Sh3^+/-^* +LPS vs. WT*+*LPS, only 4 genes in total (two are protein-coding genes) were identified (**Fig. 3E-F** highlighted with red underline). Particularly, *Acr* is a *Shank3* subordinate adjacent gene, whose expression increased when *Shank3* is deleted ^46^. To further characterize the function of DEGs from RNA-seq, we performed the pathway analysis using the QIAGEN Ingenuity Pathway Analysis (IPA) ^47^. We identified more activated inflammation-associated pathways in response to LPS challenges in both *Sh3^+/-^* and WT (**Supplementary Fig. 4 and 5)**. In contrast, inhibited neurotransmission-related pathways including NMDA receptor and calcium signaling pathways were only observed in LPS-treated *Sh3^+/-^* but not in LPS-treated WT (**Supplementary Fig. 4 and 5)**. To understand the distinct molecular pathways in LPS-treated *Sh3^+/-^* mice, we delved into the pathway analysis from *Sh3^+/-^*+LPS vs. WT+LPS (**Supplementary Fig. 6**). The top five activated pathways were all related to inflammatory processes. The top two inhibited pathways, cardiac conduction and CREB signaling in neurons, represent inhibited neuron signal transmission and neuronal functions. The other three inhibited pathways, HIF1α signaling, eNOS signaling and phagosome formation pathways, represent increased neuronal vulnerability to stress and inflammatory stimuli. In addition to the top five pathways with highest z score value, we noticed that most of the upregulated signaling pathways in response to LPS were all inflammation-related, including cytokine storm pathway and neuroinflammation pathway (**Fig. 3G**). In contrast, the downregulated pathways in LPS- treated *Sh3^+/-^* mice were neuronal and synaptic function-related, including potassium channel pathway, glutamatergic signaling pathway, acetylcholine signaling pathway, and gap junction signaling pathway (**Fig. 3G**). Overall, our RNA-seq results suggested that a combination of exacerbated neuroinflammatory responses and impaired neuronal functions is attributed to the LPS-induced behavioral regression in *Sh3^+/-^* mice.

**Figure 3.**
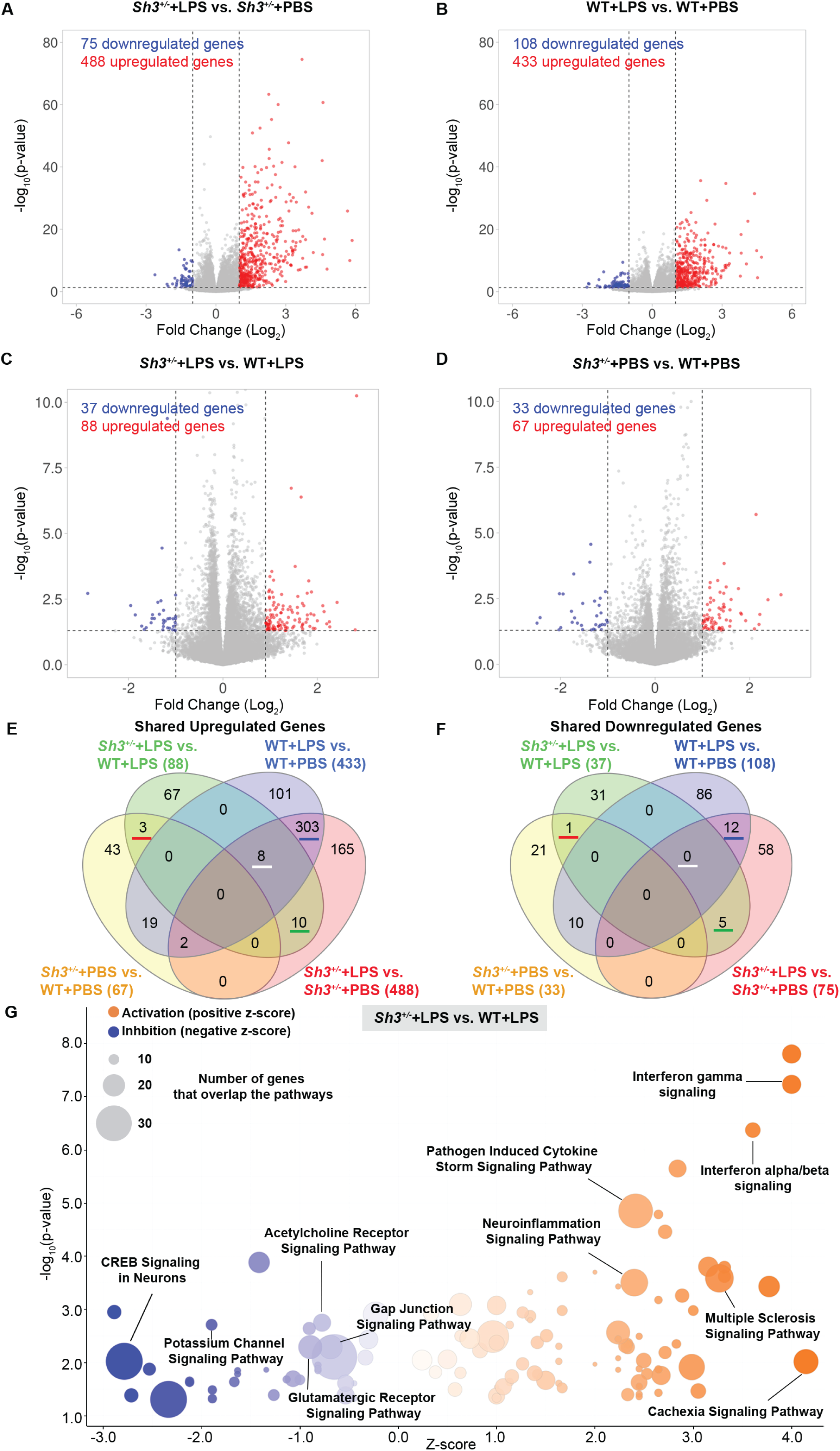
Bulk RNA sequencing revealed distinct molecular mechanism underlying LPS effect on *Sh3^+/-^* mice. **A-D. The volcano plot shows differentially expressed genes (DEGs) distribution**. DEGs were identified in four comparisons: *Sh3^+/-^+*LPS vs *Sh3^+/-^+*PBS (**A**), WT+LPS vs. WT+PBS (**B**), *Sh3^+/-^+*LPS vs. WT+LPS (**C**), *Sh3^+/-^+*PBS vs. WT*+*PBS (**D**). Upregulated genes are indicated in red color, downregulated genes are indicated in blue color and DEGs below threshold are indicated in grey color. **E-F. Venn diagrams show shared upregulated and downregulated genes among four comparisons.** Shared upregulated (**E**) and downregulated genes (**F**) numbers are shown within the diagrams. Comparison of *Sh3^+/-^+*LPS vs *Sh3^+/-^+*PBS is red diagram, WT+LPS vs. WT+PBS is blue diagram, *Sh3^+/-^+*LPS vs. WT+LPS is green diagram, and *Sh3^+/-^+*PBS vs. WT+PBS is yellow diagram. G. Bubble chart shows functions of DEGs of *Sh3^+/-^*+LPS vs. WT+LPS comparison. Volcano plot of the negative log of p-value (Y-axis) vs. the z-score (X-axis). *p*-value was calculated using the right-tailed Fisher’s Exact Test. A negative log of p-value cutoff of 1.3 (*p* < 0.05) was used to identify significantly changed functional pathways. A z-score range cutoff of ±0.1 was used to identify the activation or inhibition state of a canonical pathway. Some names of pathways are annotated in the chart. Activated pathways are orange bubbles and inhibited pathways are blue bubbles. The size of the bubbles reflects the number of genes overlapped with the pathway database.

**Table 1.** Eight Upregulated DEGs shared among *Sh3^+/-^*+LPS vs. WT+LPS, *Sh3^+/-^*+LPS vs. *Sh3^+/-^*+PBS and WT+LPS vs. WT+PB. S.

| Gene name | Protein function |
| --- | --- |
| *Bcl3* | BCL3 is an autonomous determinant of memory/terminal effector cell balance during CD8<sup>+</sup> T cell differentiation in response to acute viral infection <sup>58</sup>. |
| *Ccl3* | CCL3 as a cytokine contributes to progressive tissue damage and functional impairment during secondary injury after spinal cord injury <sup>59</sup>. |
| *Cxcl10* | CXCL10 as a lymphocyte chemoattractant contributes to the intraparenchymal trafficking of lymphocytes during acute CNS inflammation <sup>60</sup>. |
| *Gbp5* | GBP5 as a GTPase participates in the regulation of immunity and neuroinflammation <sup>61</sup>. |
| *Ifitm6* | IFITM6 encodes cell surface proteins that modulate cell-cell adhesion and cell differentiation, is induced in various macrophages, and increases expression 24 hours after LPS <sup>62</sup>. |
| *Il1b* | IL-1β can be acutely induced in the brain following peripheral bacterial infections, and microglia facilitate exaggerated IL-1β production upon exposure to acute systemic inflammation induced by bacterial LPS <sup>63</sup>. |
| *Steap4* | STEAP4 as a key effector molecule contributes to the pathogenesis of neuroinflammation in experimental autoimmune encephalomyelitis <sup>64</sup>. |
| *Tlr2* | Toll-like receptors (TLRs) are fundamental sensor molecules of the host innate immune system, which detect conserved molecular signatures of a wide range of microbial pathogens and initiate innate immune responses via distinct signaling pathways <sup>65</sup>. |

**Table 2.** Ten upregulated DEGs only shared between *Sh3^+/-^*+LPS vs. WT+LPS and *Sh3^+/-^* +LPS vs. *Sh3^+/-^*+PBS, not include in WT+LPS vs. WT+PBS. .

| Gene name | Protein function |
| --- | --- |
| *Alms1-ps1* | ALMS1, centrosome and basal body associated, pseudogene 1. |
| *Atf3* | ATF3 as a transcription factor responds to and exert dual effects on inflammatory responses and in involved in neuroinflammatory response <sup>66</sup>. |
| *Ccl12* | CCL12 as a chemokine expressed by microglia is involved in inflammation <sup>67</sup>. |
| *Fas* | Fas/FasL pathway plays a key role in immune homeostasis and leads to excessive neuroinflammation during excessive neuroinflammation <sup>68</sup>. |
| *Gm11681* | long non-coding RNA. |
| *Hspa1a* | HSPA1A and HSPA1B are two subtypes of HSPA1 proteins. HSPA1 is a member of the stress-inducible 70-kDa heat shock protein (HSP70) family, which plays a critical role in the cellular stress response <sup>69,70</sup>. |
| *Hspa1b* |  |
| *Hspb1* |  |
| *Oas1g* | OAS1G is an important murine antiviral response factor, function through activating RNase L to degrade viral RNA in turn inhibiting viral replication and propagation <sup>71</sup>. |
| *Prok2* | PROK2 is a secreted protein involved in the pathogenesis of several acute and chronic neurological diseases. It shows that PROK2 mediates neuronal cell deaths in traumatic brain injury via ferroptosis <sup>72</sup>. |

**Table 3.**
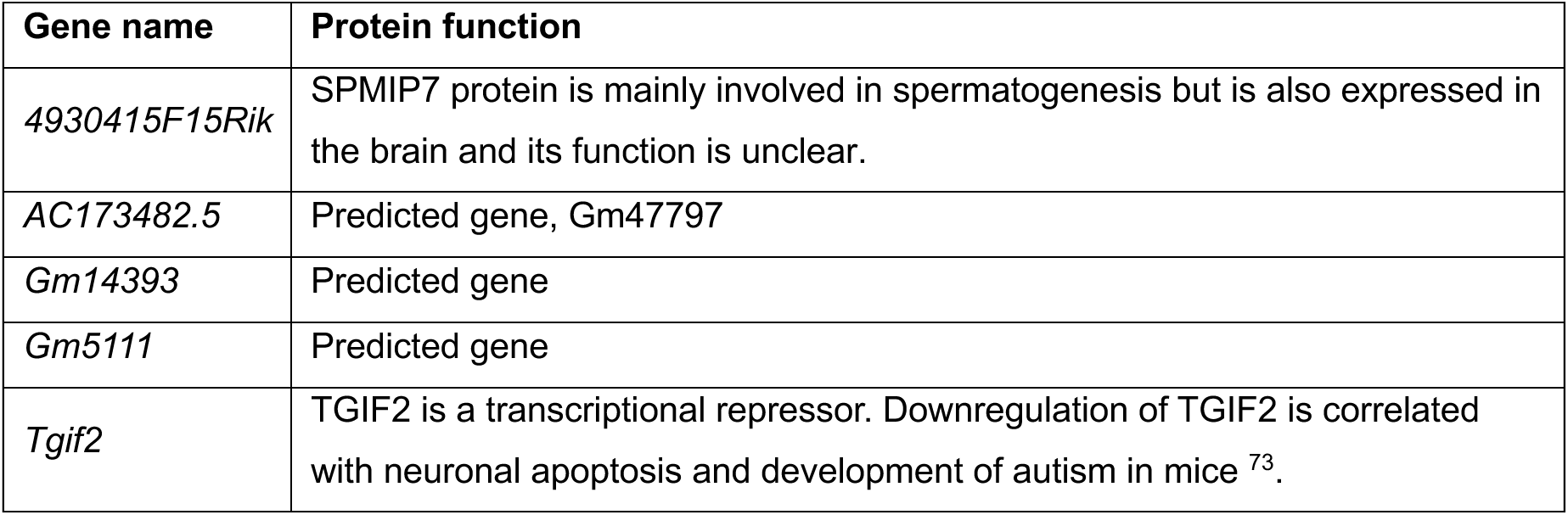
Five downregulated DEGs only shared between *Sh3^+/-^*+LPS vs. WT+LPS and *Sh3^+/-^*+LPS vs. *Sh3^+/-^*+PBS, not include in WT+LPS vs. WT+PBS. .

| Gene name | Protein function |
| --- | --- |
| *4930415F15Rik* | SPMIP7 protein is mainly involved in spermatogenesis but is also expressed in the brain and its function is unclear. |
| *AC173482.5* | Predicted gene, Gm47797 |
| *Gm14393* | Predicted gene |
| *Gm5111* | Predicted gene |
| *Tgif2* | TGIF2 is a transcriptional repressor. Downregulation of TGIF2 is correlated with neuronal apoptosis and development of autism in mice <sup>73</sup>. |

### Neuroinflammation was increased in *Sh3^+/-^* mice

To investigate how LPS leads to aggravated neuroinflammation in *Sh3^+/-^*brains, we first measured the RNA expression of the pro-inflammatory cytokines *Il1b* and chemokines *Cxcl10* using RT-qPCR. *Il1b* and *Cxcl10* have been identified as common upregulated genes across all LPS-treated groups in RNA-seq (**Table 1**). RT-qPCR confirmed that LPS significantly increased *Il1b* and *Cxcl10* expression in both WT and *Sh3^+/-^* mice compared to PBS groups. Although no significant difference in *Il1b* and *Cxcl10* expression was found between *Sh3^+/-^*+PBS vs. WT+PBS, *Sh3^+/-^*+LPS showed significantly increased expression compared to WT+LPS group (**Fig. 4A-B**). A previous study showed that LPS treatment upregulates microglia marker gene *Cx3cr1* while downregulates microglia homeostasis gene *P2ry12* expression ^48^. Therefore, we examined the expression of microglia markers *Cx3cr1* and *P2ry12* using RT-qPCR. We found that, as expected, LPS significantly increased *Cx3cr1* expression and decreased *P2ry12* expression in both WT and *Sh3^+/-^*mice compared to PBS groups (**Fig. 4C-D**). However, *Cx3cr1* expression was comparable between *Sh3^+/-^*+PBS and WT+PBS but significantly increased in *Sh3^+/-^*+LPS compared to WT+LPS group (**Fig. 4C**). In contrast, *P2ry12* expression was significantly higher in *Sh3^+/-^*+PBS than in WT+PBS and higher in *Sh3^+/-^*+LPS than in WT+LPS (**Fig. 4D**).

**Figure 4.**
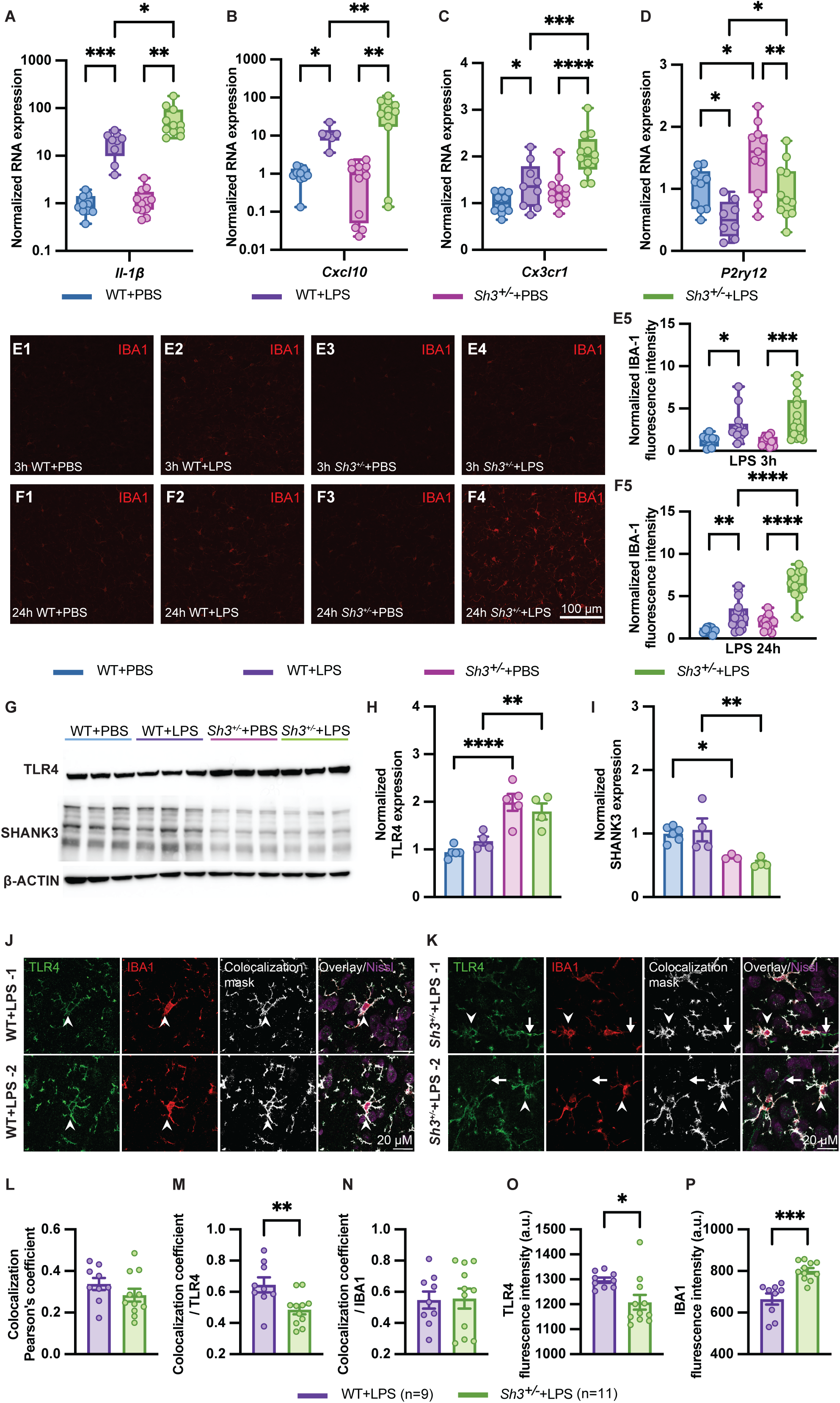
Neuroinflammation increased in *Sh3^+/-^* mice compared to WT mice in response to LPS. **A-D. Differential expression of neuroinflammatory marker genes *Il1b*, *Cxcl10, Cx3cr1* and *P2ry12***. At 24 hours after LPS injection, forebrain RNA was extracted, and RT-qPCR was performed. Target gene expression was normalized to the expression of the housekeeping gene GAPDH. One-way ANOVA was performed to compare gene expression in four groups (*Il1b*: F= 14.93, *p* = 0.0004, *Cxcl10*: F= 16.73, *p* = 0.0002, *Cx3cr1*: F= 14.8, *p* < 0.0001, *P2ry12*: F= 8.442, *p* = 0.0002). As expected, compared to PBS treatment, LPS significantly increased expression of *Il1b, Cxcl10*, and *Cx3cr1*, and decreased expression of *P2ry12* in both WT and *Sh3^+/-^* mice (*Il1b*: WT+PBS (n=10) vs. WT+LPS (n=9), *p* = 0.0005, *Cxcl10*: WT+PBS (n=10) vs. WT+LPS (n=6), *p* = 0.0105, *Cx3cr1*: WT+PBS (n=10) vs. WT+LPS (n=9), *p* = 0.0388, *P2ry12*: WT+PBS (n=10) vs. WT+LPS (n=8), *p* = 0.0241; *Il1b*: *Sh3^+/-^* +PBS (n=12) vs. *Sh3^+/-^* +LPS (n=11), *p* = 0.002, *Cxcl10*: *Sh3^+/-^* +PBS (n=12) vs. *Sh3^+/-^*+LPS (n=11), *p* = 0.0015, *Cx3cr1*: *Sh3^+/-^* +PBS (n=12) vs. *Sh3^+/-^* +LPS (n=12), *p* < 0.0001, *P2ry12*: *Sh3^+/-^* +PBS (n=11) vs. *Sh3^+/-^* +LPS (n=11), *p* = 0.0046). Significant increase of *Il1b, Cxcl10*, *Cx3cr1* was observed in *Sh3^+/-^* +LPS vs. WT+LPS (*Il1b*: *p* = 0.0166, *Cxcl10*: *p* = 0.0076, *Cx3cr1*: *p* = 0.0004). In contrast, significantly increased expression of *P2ry12* was observed in *Sh3^+/^*^-^+PBS vs. WT+PBS (*p* = 0.0102) and *Sh3^+/-^*+LPS vs. WT+LPS (*p* = 0.036). **E-F. Increased immunohistochemistry fluorescence intensity of IBA1 in *Sh3^+/-^* +LPS group**. Mouse brain slices were collected at 3 hours and 24 hours after injection, respectively. Microglia activation was labeled with IBA1 antibody (**E1-E4, F1-F4**). Quantitative fluorescence intensity of IBA1 at cortex regions from WT+LPS, *Sh3^+/^*^-^+PBS, and *Sh3^+/^*^-^+LPS groups were normalized to WT+PBS group and then compared (Kruskal-Wallis test, E5. *p* = 0.0002; F5. *p* < 0.0001). LPS increased microglia activation in both WT and *Sh3^+/^*^-^ mice at both 3 hours and 24 hours after injection compared to PBS groups (3 hours: WT+PBS (n=10) vs. WT+LPS (n=13), *p* = 0.0067, *Sh3^+/-^* +PBS (n=12) vs. *Sh3^+/-^* +LPS (n=11), *p* = 0.0004. 24 hours: WT+PBS (n=9) vs. WT+LPS (n=16), *p* = 0.0053, *Sh3^+/-^* +PBS (n=15) vs. *Sh3^+/-^* +LPS (n=12), *p* < 0.0001). Moreover, LPS induced significant overactivation of microglia in *Sh3^+/^*^-^ mice at 24 hours after injection compared to WT mice, not 3 hours (WT+LPS vs. *Sh3^+/-^* +LPS, 3 hours, *p* = 0.6204, 24 hours, *p* = 0.0017). **G-I. Increased TLR4 and decreased SHANK3 protein expression in S*h3^+/-^* mice**. LPS receptor TLR4 and SHANK3 expression were examined in whole-cell lysates using western blot at 24 hours after injection. Target protein expression was normalized to β-ACTIN expression (**G**). Quantitative protein expression from WT+LPS, *Sh3^+/^*^-^+PBS, and *Sh3^+/^*^-^+LPS were normalized to WT+PBS group and one-way ANOVA was performed (TLR4, F = 14.38, *p* = 0.0001, SHANK3, F = 7.681, *p* = 0.0033). Significantly elevated TLR4 expression was observed in *Sh3^+/^*^-^ mice compared to WT mice (**H**, WT+PBS vs. *Sh3^+/-^* +PBS, *p* < 0.0001; WT+LPS vs. *Sh3^+/-^* +LPS, *p* = 0.0081). As expected, significantly reduced SHANK3 expression was detected in *Sh3^+/^*^-^ mice (**I**, WT+PBS vs. *Sh3^+/-^*+PBS, *p* = 0.0172; WT+LPS vs. *Sh3^+/-^* +LPS, *p* = 0.002). J-P. Colocalization analysis of TLR4/IBA1 between WT+LPS vs. *Sh3^+/-^* +LPS group. Mouse brain slices were collected at 24 hours after LPS injection in WT and *Sh3^+/^*^-^ mice. IBA1 and TLR4 antibody were used for immunocytochemistry. Colocalization of IBA1 and TLR4 at cortex regions were analyzed and compared (**J-K**). Unpaired t-test was conducted. No difference in colocalization Pearson’s coefficient between WT+LPS vs. *Sh3^+/-^* +LPS was observed (**L**, two-tailed *p* = 0.2309). *Sh3^+/-^* showed smaller normalized colocalization by TLR4 (**M**, two-tailed *p* = 0.0093) but similar normalized colocalization by IBA1 (**N**, two-tailed *p* = 0.9141) compared to WT. *Sh3^+/-^* showed decreased TLR4 cellular fluorescence intensity (**O**, two-tailed *p* = 0.0183) but increased IBA1 cellular fluorescence intensity (**P**, two-tailed *p* = 0.0001) than WT. * *p* < 0.05, ** *p* < 0.005, *** *p* < 0.0005, **** *p* < 0.0001.

We further examined microglia activation by examining the expression of IBA-1, a marker widely used for the assessment of microglia activation state ^49^ (**Fig. 4E-F**). Compared to PBS treatment, LPS increased IBA-1 staining intensity significantly at 3 hours and 24 hours after LPS treatment in both WT and *Sh3*^+/-^ mice. However, *Sh3*^+/-^ +LPS showed significantly higher microglia activation than WT+LPS group at 24 hours but not at 3 hours post-injection. These results indicated that the prolonged activation of microglia contributed to increase of neuroinflammatory responses in *Sh3*^+/-^ mice.

Next, to understand the mechanism underlying the LPS-induced microglia overactivation in *Sh3*^+/-^ mice, we analyzed the LPS binding receptor toll-like receptor-4 (TLR-4) expression in WT+LPS and *Sh3^+/-^*+LPS groups at 24 hours post-injection ^50^. We found that TLR4 expression was significantly higher in *Sh3^+/-^* than WT with either PBS or LPS treatment (**Fig. 4G-H**). Additionally, we measured the SHANK3 protein expression and found that either PBS- or LPS-treated *Sh3^+/-^* groups showed a reduction in SHANK3 protein expression compared to WT groups as expected, but no changes in SHANK3 expression were observed after LPS treatment (**Fig. 4G, I**). TLR4 is known to associate with LPS-induced microglia activation^51–54^. To investigate whether observed TLR4 augmentation directly facilitates microglia overactivation in LPS treated *Sh3^+/-^* mice, we analyzed colocalization between TLR4 and IBA1 (**Fig. 4J-P**). We found that TLR4 and IBA1 expression were well-colocalized in WT, but a small fraction of TLR4 signals in *Sh3^+/-^*mice were not colocalized with IBA1 (**Fig 4J-K**). Although colocalization Pearson’s coefficient was similar between WT+LPS and *Sh3^+/-^*+LPS group, colocalization coefficient normalized by TLR4 was lower in *Sh3^+/-^* mice and the colocalization coefficient normalized by IBA1 was similar (**Fig. 4L-N**). Interestingly, cellular immunofluorescence intensity of TLR4 in *Sh3^+/-^* mice is slightly lower than WT (**Fig. 4P**). In addition, cellular immunofluorescence intensity of IBA1 was significantly higher in *Sh3^+/-^* brains (**Fig. 4O**). Given that TLR4 was expressed in many different cell types including astrocytes, neutrophils, and brain endothelial cells ^55,56^, these findings suggested that elevated TLR4 expression in other cell types might contribute to microglia activation in *Sh3^+/-^* mice.

### Microglia activation affected synaptic structure in *Sh3^+/-^* mice

To uncover how microglia overactivation leads to behavioral changes in *Sh3^+/-^* mice, we examined microglia engulfment and synaptic structures. We compared lysosome engulfment (lysosome marker CD68 occupancy) and synapse engulfment (presynaptic marker vGluT1 occupancy) between WT+LPS and *Sh3^+/-^*+LPS brains at 24 hours post-injection. While CD68 occupancy was comparable between the two groups, vGluT1 occupancy — both within CD68-positive lysosomal puncta in microglia (CD68^+^IBA1^+^) and within the microglial cytoplasm (IBA1^+^) — was significantly higher in *Sh3^+/−^* mice compared to WT mice (**Fig. 5A-D**). These findings suggest that microglial activation is associated with increased phagocytosis of synaptic materials in the *Sh3^+/−^*+LPS group. To determine whether increased synaptic engulfment leads to synapse dismantling in the *Sh3^+/−^* brain, we examined the colocalization of the presynaptic marker vGluT1 with the postsynaptic density marker PSD95 at 24 hours following LPS injection. In WT mice, vGluT1 and PSD95 puncta were well colocalized (**Fig. 5E**, arrowhead). In contrast, *Sh3^+/−^* mice displayed discrete PSD95 clusters that did not colocalize with vGluT1 puncta (**Fig. 5E**, arrow). Quantitative analysis revealed that the *Sh3^+/−^*+LPS group showed a significantly lower Pearson’s correlation coefficient for colocalization, a reduced colocalization coefficient normalized by PSD95, and decreased overall synaptic puncta density compared to the WT+LPS group (**Fig. 5F-H**).

**Figure 5.**
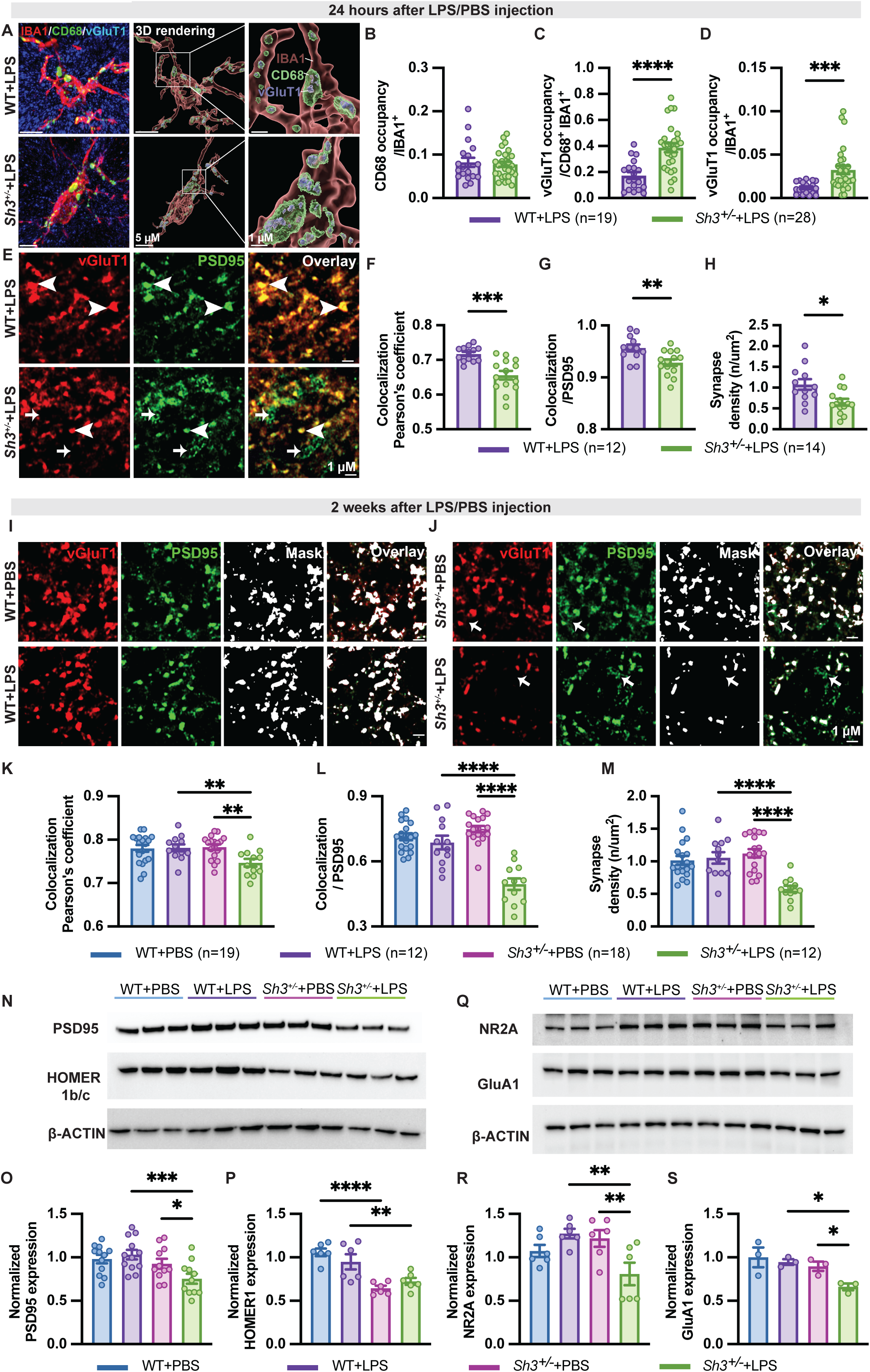
Microglia activation disrupted synaptic structure in *Sh3^+/-^* mice compared to WT mice in response to LPS. **A-D. Increased vGluT1 engulfment in *Sh3^+/-^* mice at 24 hours after LPS injection**. A. Representative images of maximum projection of z-stack imaging of IBA1/CD68/vGluT1 co-staining (left), 3D rendering masked by IBA1 and CD68 (center), and enlarged images of 3D rendering (right). Total volumes of masked vGluT1 and CD68 were calculated and unpaired t-test was conducted. No significant difference of IBA1 masked CD68 occupancy was detected (**B.** two-tailed *p* = 0.7149). IBA1 and CD68 double masked vGluT1 occupancy and IBA1 masked vGluT1 occupancy was higher in *Sh3^+/-^* mice than WT mice (**C.** two-tailed *p* < 0.0001, **D.** two-tailed *p* = 0.0003). **E-H. Reduced synapses in *Sh3^+/-^* mice at 24 hours after LPS injection**. E. Representative images of super-resolution imaging of vGluT1/PSD95 co-staining at 24 hours post-injection. Colocalized puncta of vGluT1and PSD95 were indicated by arrowheads and were counted as synapse puncta. Non-colocalized puncta of PSD95 were indicated by arrows. Colocalization of vGluT1and PSD95 was analyzed, and number of colocalized puncta were counted as synapse density. Unpaired t-test was conducted. *Sh3^+/-^* mice showed a lower colocalization Pearson’s coefficient (**F**, two-tailed *p* = 0.0002), a reduced normalized colocalization coefficient by PSD95 (**G**, two-tailed *p* = 0.0056) and decreased synapse density (**H**, two-tailed *p* = 0.0107) compared to WT mice. **I-M. Reduced synapses in *Sh3^+/-^* mice at 2 weeks after LPS injection**. I-J. Representative images of super-resolution imaging of vGluT1/PSD95 co-staining at 2 weeks post-injection among four groups, WT+PBS, WT+LPS, *Sh3^+/-^*+PBS, and *Sh3^+/-^*+LPS. Mask was generated based on the colocalized puncta of vGluT1and PSD95. Non- colocalized puncta of PSD95 were indicated by arrows in **J**. One-way ANOVA was conducted for comparing colocalization Pearson’s coefficient (**K.** F = 4.254, *p* = 0.0088), colocalization coefficient by PSD95 (**L.** F = 26.51, *p* < 0.0001), and synapse density (**M.** F = 10.94, *p* < 0.0001). *Sh3^+/-^*+LPS group showed a lower colocalization Pearson’s coefficient than WT+LPS (*p* = 0.0067) or *Sh3^+/-^*+PBS groups (*p* = 0.0022), a reduced normalized colocalization coefficient by PSD95 than WT+LPS (*p* < 0.0001) or *Sh3^+/-^*+PBS groups (*p* < 0.0001), and decreased synapse density than WT+LPS (*p* < 0.0001) or *Sh3^+/-^*+PBS groups (*p* < 0.0001). **N-S. Reduced PSD95, HOMER 1b/c, NR2A and GluA1 expression in *Sh3^+/-^* mice at 2 weeks after LPS injection.** Postsynaptic scaffold proteins PSD95 and HOMER 1b/c (**N**), and excitatory receptors NMDA subunit NR2A and AMPA subunit GluA1 expression (**Q**) were examined in forebrain PSD fractions using western blot at 2 weeks after injection. Target protein expression was normalized to β-ACTIN expression. Quantitative protein expression from WT+LPS, *Sh3^+/^*^-^+PBS, and *Sh3^+/^*^-^+LPS were normalized to WT+PBS group and one-way ANOVA was conducted (PSD95, F = 4.763, *p* = 0.006, HOMER 1b/c, F = 12.28, *p* < 0.0001, NR2A, F = 5.06, *p* = 0.0091, GluA1, F = 4.951, *p* = 0.0313). **O.** Significantly decreased PSD95 expression was detected in *Sh3^+/-^* mice not WT mice after LPS treatment (*Sh3^+/-^*+LPS (n=11) vs. WT+LPS (n=12), *p* = 0.0009; *Sh3^+/-^*+LPS (n=11) vs. *Sh3^+/-^*+PBS (n=11), *p* = 0.0354). No significant change in PSD95 was detected between WT+PBS (n=12) vs. WT+LPS (*p* = 0.5153) or WT+PBS vs. *Sh3^+/-^*+PBS (*p* = 0.4845). **P.** Significantly reduced HOMER 1b/c expression was observed in *Sh3^+/^*^-^ mice compared to WT mice (WT+PBS (n=6) vs. *Sh3^+/-^* +PBS (n=6), *p* < 0.0001; WT+LPS (n=6) vs. *Sh3^+/-^*+LPS (n=6), *p* = 0.0088). No significant change of HOMER 1b/c was detected after LPS treatment (WT+PBS vs. WT+LPS (*p* = 0.1781) or *Sh3^+/-^*+PBS vs. *Sh3^+/-^*+LPS (*p* = 0.318)). **R.** *Sh3^+/-^*+LPS group (n=6) showed significant decreased NR2A expression compared to both the *Sh3^+/-^*+PBS group (n=6) (*p* = 0.0053) and the WT+LPS group (n=6) (*p* = 0.002). No significant change of NR2A was observed between WT+PBS (n=6) vs. WT+LPS (*p* = 0.1404) or WT+PBS vs. *Sh3^+/-^*+PBS (*p* = 0.2827). **S.** *Sh3^+/-^*+LPS group (n=3) showed significantly reduced GluA1 expression compared to both the *Sh3^+/-^* +PBS group (n=3) (*p* = 0.0378) and the WT+LPS group (n=3) (*p* = 0.0169). No significant change in GluA1 was found between WT+PBS (n=3) vs. WT+LPS (*p* = 0.5898) or WT+PBS vs. *Sh3^+/-^*+PBS (*p* = 0.3102). * *p* < 0.05, ** *p* < 0.005, *** *p* < 0.0005, **** *p* < 0.0001.

Next, we compared the colocalization of vGluT1 and PSD95 among four groups — WT+PBS, WT+LPS, *Sh3^+/−^*+PBS, and *Sh3^+/−^*+LPS — at two weeks after injection. In both WT+PBS and WT+LPS groups, vGluT1 and PSD95 puncta remained closely aligned (**Fig. 5I**). The *Sh3^+/−^*+PBS group also exhibited substantial colocalization of vGluT1 and PSD95; however, some displaced PSD95 fragments were observed, which became more pronounced in the *Sh3^+/−^*+LPS group (**Fig. 5J**, arrow). Consistent with the findings at 24 hours post-injection, the *Sh3^+/−^*+LPS group showed a significantly lower Pearson’s correlation coefficient for colocalization, a reduced colocalization coefficient normalized by PSD95, and decreased synapse density compared to the other groups (**Fig. 5K-M**). These results support the conclusion that microglial overactivation enhances synapse engulfment and contributes to synaptic structure disorganization in *Sh3^+/−^* mice in response to LPS-induced inflammation.

To investigate whether disrupted synaptic structure caused PSD protein reduction and synaptic dysfunction in *Sh3^+/-^* mice, we explored excitatory postsynaptic scaffold proteins PSD95 and HOMER 1b/c expression (**Fig. 5N**). No significant change of PSD95 was observed between *Sh3^+/-^*+PBS vs. WT+PBS. However, PSD95 was significantly decreased in the *Sh3^+/-^*+LPS compared to both the WT+LPS group (∼28% reduction) and the *Sh3^+/-^*+PBS group (∼17% reduction), suggesting that LPS treatment reduced the PSD95 protein level in *Sh3^+/-^* mice not in WT mice (**Fig. 5O**). In contrast, HOMER 1b/c expression was significantly decreased in *Sh3^+/-^*+PBS vs. WT+PBS (∼41% reduction) and in *Sh3^+/-^*+LPS vs. WT+LPS (∼23% reduction) but showed no significant change between *Sh3^+/-^*+LPS vs. *Sh3^+/-^*+PBS. It indicated that LPS treatment did not further reduce HOMER expression (**Fig. 5P**). PSD95 is critical for excitatory synapse function through its interactions with NMDA and AMPA receptors^57^. We examined whether the change of PSD95 may lead to the change of the expression of major subunits of NMDA receptor subunit NR2A and the AMPA receptor subunit GluA1 associated with LPS treatment (**Fig. 5Q**). We found that NR2A expression was reduced in the *Sh3^+/−^*+LPS group compared to both the WT+LPS group and the *Sh3^+/−^*+PBS group (**Fig. 5R**). The GluA1 expression was significantly reduced in *Sh3^+/−^*+LPS mice compared to both the WT+LPS group and *Sh3^+/−^*+PBS group **(Fig. 5S**). Taken together, these results suggested a mechanistic convergence between LPS-induced neuroimmune responses, and the synaptic disruptions known to arise from Shank3 deficiency.

## Discussion

In this study, we successfully established a preclinical model and experimental protocol to assess the role of neuroinflammation induced by LPS in *Sh3^+/-^* mice (**Fig. 6**). We demonstrated that LPS challenge modified the penetrance of motor function deficits, anxiety-like behavior, and repetitive behavior but did not alter social preference in *Sh3^+/-^*mice. More interestingly, the motor and emotion specific behavioral changes are reminiscent of catatonia features that are often observed in humans with *SHANK3* deficiency and idiopathic ASD ^74–76^. Additionally, our extensive comparative transcriptome analysis among the different groups validated the experimental protocol of LPS induction and revealed distinct underlying molecular mechanisms. The significantly upregulated genes ubiquitously associated with inflammatory pathways in LPS treated groups indicated the effectiveness of LPS induction. The downregulated genes in multiple pathways associated with neuronal and synaptic functions strongly support a potential causal link between LPS treatment and increased penetrance of abnormal behaviors in *Sh3^+/-^* mice. Moreover, increased microglia activation and reduced synaptic function from LPS treatment were further validated by molecular experiments. Finally, the partial reversal of motor impairment in LPS treated *Sh3^+/-^* mice through the anti-inflammation treatment with MFA further supports this causality. Notably, MFA treatment did not restore LPS-treated WT and *Sh3^+/-^*mice performance in the accelerating speed rotarod. These findings suggest that the mechanisms underlying inflammation-induced motor impairment are different between steady speed and accelerating speed rotarod tests ^77^. Additionally, one-week MFA treatment was more effective in female *Sh3^+/-^* mice than male mice (**Fig. 2D-E**) and LPS-treated male *Sh3^+/-^* mice performance was worse than female ones (**Supplementary Fig. 1A-B**), indicating potential sex difference in motor function^78^ and neuroinflammation process^79^.

**Figure 6.**
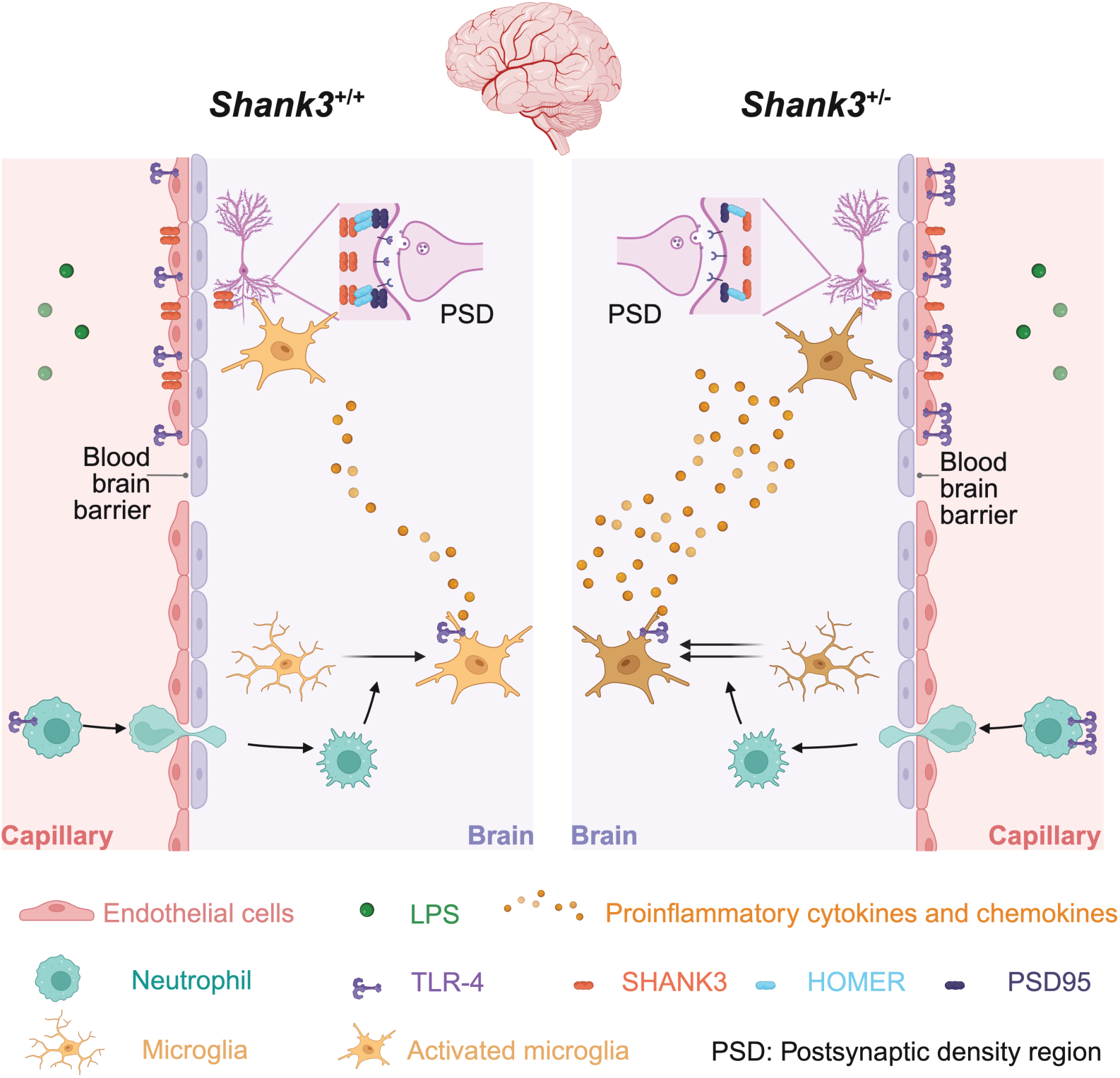
Summary of increased neuroinflammation susceptibility in *Sh3^+/-^* mice. The schematic drawing proposes a model of increased neuroinflammation and altered behavioral phenotypes in *Shank3* haploinsufficiency mouse after LPS treatment. Generally speaking, LPS evokes neuroinflammation through three distinct pathways, 1) LPS binds to TLR4 expressed on neutrophils, promoting neutrophil infiltration through blood-brain barrier and activation of microglia cells; 2) LPS binds to TLR4 expressed in cerebral endothelial cells located in blood-brain barrier, compromising the integrity of blood-brain barrier and increasing permeability of additional immune components entering the brain; 3) LPS that penetrates blood-brain barrier binds to TLR4 expressed in microglia, activating microglia and triggering proinflammatory cytokines release. In *Sh3^+/-^*brain vs. WT brain, we observed excessive proinflammatory gene expression, proinflammatory cytokines, overactivation of microglia and increased expression of TLR4 at 24 hours after LPS injection. It suggests that LPS might exacerbate neuroinflammation responses through above routes in *Sh3^+/-^* mice. Microglia activation can sculpt synapses and brain circuits and change animal behavioral phenotypes long-term. In *Sh3^+/-^* brain vs. WT brain, we observed elevated microglia engulfment, reduced synapse, and decreased synaptic transmission gene expression at 24 hours after LPS injection, suggesting increased synapse elimination in *Sh3^+/-^* mice. At 2 weeks after LPS injection, we observed reduced synapse density, decreased PSD95 expression and lower expression of excitatory receptor expression. It indicates that neuroinflammation-induced synaptic dysfunction was associated with behavioral changes found in *Sh3^+/-^*mice.

The behavior impairments induced by LPS in *Sh3^+/-^* mice could attribute to the dysfunction of neuronal cells resulted from neuroinflammation. In the previous study that our lab contributed, LPS challenge increased the excitability in D1 but not D2 medium spiny neurons using whole-cell patch clamp recordings and decreased neuron calcium transients in nucleus accumbens in response to social stimuli using *in vivo* Ca^2+^ imaging^33^. Our data showed that increased neuroinflammation downregulated genes associated with neuronal and synaptic functions in *Sh3^+/-^* mice compared to WT mice and reduced synaptic protein PSD95 in *Sh3^+/-^* mice.

LPS-induced neuroinflammation could also alter the function of non-neuronal cells in *Sh3^+/-^* mice. Our findings of augmented TLR4 expression and microglia activation associated with LPS treatment suggested that microglia are primarily involved in more significant neuroinflammatory responses in *Sh3^+/-^* mice. Consistent with the findings in many reports in literature supporting that TLR4 mediates microglial activation in brains ^51-^_54_, our findings also support that the upregulation of TLR4 contributes to the observed microglial activation following LPS administration. The direct causality between upregulation of TLR4 and microglia activation due to LPS induction has been demonstrated in a recent study^80^. Our observations are consistent with this finding and support the augmented TLR4 as the cause of microglia activation in SHANK3 deficiency mice. Further analysis of TLR4 and IBA1 cellular colocalization indicated that additional cell types might also be involved in LPS-TLR4 (ligand-receptor) mediated neuroinflammation. Cellular fluorescence intensity of TLR4 in fixed brain cortex regions decreased while total protein level of TLR4 in freshly prepared forebrain tissue increased in *Sh3^+/-^* mice than WT. This inconsistence implied that changes of TLR4 in *Sh3^+/-^* might be brain region specific and cell type specific, which warranted a future study on the TLR4 transduction signaling pathways in *Shank3* haploinsufficiency mouse model.

Moreover, we uncovered a mechanistic convergence between LPS-induced neuroimmune responses, and the synaptic disruptions known to arise from Shank3 deficiency. We discovered increased microglia engulfment and disrupted synaptic structure in *Sh3^+/-^* mice. Such dysfunction sustained until 2 weeks later when we identified LPS-triggered behavioral changes only in *Sh3^+/-^* mice instead of WT mice. Moreover, in a previous study, we reported that the scaffold protein PSD95 expression did not change significantly while HOMER1b/c was significantly reduced in *Sh3^+/−^*mice^35^. In the current study, we replicated the finding that HOMER1b/c expression is significantly decreased in *Sh3^+/−^* mice. However, LPS treatment did not further reduce HOMER levels. We also replicated the finding that PSD95 is not changed in *Sh3^+/−^* mice. However, LPS treatment significantly reduced PSD95 expression in *Sh3^+/−^* mice not WT mice. This pattern of PSD95 changes in response to LPS, influenced by Shank3 haploinsufficiency, differs from that observed for HOMER1b/c. Although we previously reported no change of NR2A or GluA1 in *Sh3^+/-^* mice^35^, we found that LPS treatment reduced NR2A and GluA1 expression in *Sh3^+/-^*mice instead of WT mice.

Our findings represent a first step toward understanding how Shank3 haploinsufficiency is linked to neuroimmune signaling. Our previous study showed that SHANK3 deficiency in periphery nerve system contributes to the exaggerated inflammatory responses^81^. Our recently published collaborative study has shown that conditional knock-out of SHANK3 major isoform in brain endothelial cells recapitulated social behavior impairment and other synaptic phenotypes ^82^. Our findings are aligned with previous studies showing that *Shank3* deficiency exacerbates the inflammatory responses in peripheral neurons either induced by LPS ^34,81^ or by maternal immune activation during the pregnancy ^83^. Overall, our findings showed that LPS exacerbated neuroinflammation in *Sh3^+/-^*mouse brains through unique molecular mechanisms as summarized in **Fig. 6** for further investigations in future.

In humans, haploinsufficiency of *SHANK3* is highly penetrant for neurodevelopmental impairments ^84^. Approximately ∼70% of PMS and *SHANK3* deficiency patients met the diagnostic criteria for ASD. The comorbidity of intellectual disability, anxiety, and epilepsy are highly penetrant or common ^75^. There is an increasing recognition of behavioral regression and catatonia among young teens with *SHANK3* deficiency ^74^. However, the triggers and mechanisms underlying the behavioral regression are largely unknown. Although few clinical case reports implicated neuroinflammation as a trigger ^21,85^, our findings may offer a mechanistic explanation for the behavioral regressions observed in *SHANK3* patients after experiencing a subclinical infection responded to the immune modulator treatment. Our study may support a link between neuroinflammation and autism susceptibility in general.

Our results establish an important proof-of-concept study to dissect the link between neuroinflammation and autism susceptibility. The apparent limitations of our studies include that we have not provided definitive causality for the cell types and specific molecular mechanism directly implicated in the increased neuroinflammatory responses indued by LPS in SHANK3 haploinsufficiency mice. Future studies are warranted to examine dosage- or treatment duration-dependent effects of LPS on behavioral outcomes at different neurodevelopmental stages and in different sexes of *Shank3* deficiency mice. Despite these limitations, our results suggest that *Shank3* haploinsufficiency increases susceptibility to neuroinflammation-induced behavior impairment through distinct molecular pathways. These findings provide new insights into the role of neuroinflammation in ASD and for developing anti-inflammatory treatments as both preventive and therapeutic strategies for behavior regression in ASD that are warranted for further investigations.

## Materials and methods

### Animals

Animals were housed under standard 12 h/12 h light/dark cycles (7 AM-7 PM) with food, water ad libitum, controlled temperature and humidity in the Yale University animal facility. Experiments procedures conducted were following the Institutional Animal Care and Use Committee (IACUC) protocol at Yale University (protocol No. 2025-20271). *Shank3*^Δe4–22^ mouse line (Jax Strain #:039524) was generated in-house and had been crossed with C57BL/6J mouse line (Jax Strain #:000664) over 20 generations. Wild-type (*WT*), *Shank3*^Δe4–22^ homozygous (*Sh3^-/-^*) and *Shank3*^Δe4–22^ heterozygous mice (*Sh3^+/-^*) were obtained through heterozygotes breeding as previously described ^35^. All experiment procedures were performed within a time frame of 9 AM-5 PM. Animals of the same sex were group housed (2-5 mice per cage). Both male and female mice were used. One cage of mice with same genotype and same sex were assigned to PBS or LPS group randomly. Experimental order of different group was randomized. Mice from multiple groups for same experiments were carried out at same time. Before experiments, each mouse was cataloged with identification number in an excel sheet which included detailed information such as genotype and drug treatment. During experiments, experimenter handled mice based on its identification number and attempted to be semi-blinded. Post-processing, data handling and statistical analysis were semi-automated performed using software stated below. Mice with health problems such as runt, fight wounds, malocclusion, and eye problems, were excluded from experiments. For RNA and protein extraction, mice were deeply anesthetized with 5% isoflurane, then mouse brains were quickly collected, flash frozen in liquid nitrogen and stored in -80 °C freezer until further use. The number of animals used in each experiment was determined based on our lab prior experience and a prior power analysis was calculated using G*power statistics software^86^ (**Supplementary table 5**). Total number of mice used for each experiment was as follows: 86 mice for MFA treatment rotarod test, 52-95 mice for open field, rotarod and three-chamber behavior test, 49-66 mice for light/dark box and grooming behavior test, 15 mice for RNA sequencing, 40 mice for quantitative PCR, 24 mice for wide-field IBA1 staining, 6 mice for TLR4/IBA1 co-staining, 6 mice for microglia engulfment and vGluT1/PSD95 staining at 24 hours post-injection, 12 mice for vGluT1/PSD95 staining at 24 hours post-injection, 18 mice for whole-cell protein analysis and 24 mice for synaptic protein analysis. All experiments were replicated at least once, except that RNA sequencing was done one time.

### Drug

Three to five-week-old female and male mice were intraperitoneally injected with LPS (Sigma, L6511) in saline at 1 mg/kg or same volume of saline (PBS). To test the mefenamic acid (MFA) anti-inflammatory effect on behavior improvement, MFA (Thermofisher, AAJ6270522) dissolved in DMSO at 5 mg/kg or the same volume of DMSO (vehicle) was injected intraperitoneally to the mice with LPS injection once a day for 7 days. Before injection, freshly made stock solution of MFA in DMSO (MFA treatment) or DMSO (vehicle treatment) was diluted with 1:100 ratio in PBS to minimize DMSO toxicity.

### Behavior assay

For behavioral experiments, mice were handled for 3 days and habituated in the procedure room for 1 hour before behavioral assays. All behavioral tests were conducted once for each mouse.

Anxiety-like behaviors were tested by open field test (OFT) and light/dark box test followed previous studies^35,87^. In OFT, mouse spontaneous activity in Noldus Open Field (4 chambers, 40 (L) x 40 (W) x 30 (H) cm per chamber) over 10 minutes was recorded with EthoVision XT video tracking software (Noldus). The Noldus two-chambered light/dark box is equipped with the lid of dark chamber made by infrared translucent material. The dimensions of the compartment are one third for the dark chamber (20 (L) x 20 (W) x 20 (H) cm) and two thirds for the light chamber (20 (L) x 40 (W) x 20 (H) cm). Mice were firstly placed into the open-light chamber (∼700 lux) and given 5 min to freely explore both light and dark (0 lux) chambers. The video was recorded with EthoVision XT video tracking software. Ethovision software was used to automatically score the travel distance and duration in arena.

Repetitive behavior was tested by evaluating mouse grooming behaviors. Mouse was placed in a Noldus phenotyper (30 (L) x 30 (W) x 45 (H) cm) and recorded with EthoVision XT video tracking software. Mouse freely explored for 10 minutes and last 5 minutes of grooming behavior was hand scored.

Motor coordination was tested by rotarod (Med Associate) performance as described ^35,88^. Mice were tested in two 5 min trials with an inter-trial interval of 30 min. In accelerating session, revolutions per minute (rpm) of the turning barrel is accelerating from 4 to 40. In steady speed session, the rpm of the turning barrel is fixed at 16. Latency to fall, or to rotate off the top of the turning barrel, was measured by the timer.

Social preference was tested by a three-chamber assay as described^88^. An unfamiliar age- and sex-matched mouse was used as a social stimuli mouse in the three-chamber test and habituated under a wired enclosure for 3 days before testing. A customized chamber composed of a rectangular Plexiglas arena (42 (L) x 56.5(W) x 35.8 (H) cm) divided into three chambers (each chamber 18.5 x 42 cm), with 2 doors connecting chambers and two enclosures (wired pencil cups with metal vertical bars) located in the two outer chambers. Briefly, the test mouse was habituated in the center chamber for 5 min. Then the social stimuli mouse was placed under one of the pencil cups and 2 doors were lifted allowing the experimental mouse to freely explore the whole arena. The location of the empty and mouse enclosures, and the social stimuli mouse were changed and counterbalanced for each trial. The EthoVision XT video tracking software was used to score the duration of mouse spending in the empty chamber, the mouse chamber, proximity regions (5 cm in distance) close to the empty enclosure, and proximity regions close to the mouse enclosure.

### Bulk RNA sequencing

Total RNA was isolated from the forebrain hemisphere using Trizol (Invitrogen, 15596018) followed manufacturer protocol and then treated with DNase I (Invitrogen, AM1907) to remove contaminating DNA. RNA samples were sent to Yale Center for Genome Analysis (YCGA) for bulk RNA sequencing (RNA-seq). See details about sequencing process in Supplementary Information. After obtaining the data, low quality reads were trimmed, and adaptor contamination were removed using Trim Galore (v0.5.0). Trimmed reads were mapped to the mouse reference genome (mm10) using HISAT2 (v2.1.0) ^89^. Gene expression levels were quantified using StringTie (v1.3.3b) ^90^. Differentially expressed genes were identified using DESeq2 (v 1.22.1) ^91^. Unpaired *t*-test *p* value was used for filtering DEGs with significant differences between the two groups. Log_2_fold cutoff threshold at 1 and *p* < 0.05 was used to filter DEGs from two groups comparison (WT+LPS vs. WT+PBS, *Sh3^+/-^*+LPS vs. *Sh3^+/-^*+PBS, *Sh3^+/^*+LPS vs. WT+LPS and *Sh3^+/-^*+PBS vs. WT+PBS) for genes comparison and analysis. *Pathway analysis*: QIAGEN Ingenuity Pathway Analysis (IPA) library of canonical pathways was used to identify the pathways most significant to the data set. Molecules from the dataset meeting the criteria in which log_2_fold cutoff threshold at 0.25 and *p* < 0.05 were associated with a canonical pathway in the QIAGEN Knowledge Base were considered for the analysis. Because using log_2_fold cutoff threshold at 1 only revealed three activated pathways in *Sh3^+/^*+LPS vs. WT+LPS comparison which were all inflammation-related, so we expanded the log_2_fold cutoff threshold to 0.25. The significance of the association between the data set and the canonical pathway was measured based on: 1) a ratio of the number of molecules from the data set that map to the pathway divided by the total number of molecules that map to the canonical pathway was displayed, 2) a right-tailed Fisher’s Exact Test was used to calculate a p-value determining the probability that the association between the genes in the dataset and the canonical pathway was explained by chance alone, 3) A z-score was calculated to indicate the likelihood of activation or inhibition of that pathway. In this paper, pathways with right-tailed Fisher’s Exact Test *p* < 0.05, z-score > 0.1 was considered as activated pathways and z-score < -0.1 was considered as inhibited pathways. The volcano plot was created using VolcaNoseR ^92^. Veen diagram was plotted using InteractiVenn ^93^.

### Real-Time Quantitative Reverse Transcription PCR (RT-qPCR)

Total RNA was isolated using same protocol as RNA-seq. RNA was transcribed into cDNA using iScript cDNA Synthesis Kit (Bio-Rad, 1708890) and qPCR was performed in triplicate using iQ SYBR Green Supermix (Bio-Rad, 1708880). Gene expressions were normalized to the expression of GAPDH. Primers used to amplify cDNAs are: *Il1b,* forward, TGGACCTTCCAGGATGAGGACA, reverse, GTTCATCTCGGAGCCTGTAGTG; *Cxcl10,* forward, ATCATCCCTGCGAGCCTATCCT, reverse, GACCTTTTTTGGCTAAACGCTTTC; *Cx3cr1,* forward, GAGCATCACTGACATCTACCTCC, reverse, AGAAGGCAGTCGTGAGCTTGCA; *P2ry12* forward, CATTGACCGCTACCTGAAGACC, reverse, GCCTCCTGTTGGTGAGAATCATG; *Gapdh* forward, CAAAATGGTGAAGGTCGGTG, reverse, AATGAAGGGGTCGTTGATGG.

### Immunohistochemistry

Transcardiac perfusion was conducted at both 3 hours and 24 hours after LPS or PBS injection with 20 mL PBS then 20 mL 4% paraformaldehyde (PFA, Santa Cruz, sc-281692). Mouse brains were collected, fixed in 4% PFA for 24 hours, then changed to 30% (w/v) sucrose before sectioning. 50 μM brain sections between Bregma 1 mm to -1 mm were obtained using vibratome (Leica, VT1200S) or using cryostat (Leica, CM1860). Primary antibody including anti-rabbit-IBA1 (1:200, FUJIFILM Wako Chemicals, 019-19741), anti-mouse-TLR4 (1:100, Abcam, ab22048), anti-rat-CD68 (1:100, Abcam, ab53444), anti-mouse-vGluT1 (1:100, Synaptic Systems, 135511), anti-guineapig-vGluT1 (1:200, Synaptic Systems, 135318) and anti-mouse-PSD95 (1:100, Invitrogen, MA1-045), and secondary antibody including goat anti-mouse Alexa Fluor 488 (Abcam, ab150113), donkey anti-mouse Alexa Fluor 488 (Abcam, ab150105), donkey anti-rabbit Alexa Fluor 594 (Invitrogen, A-21207), goat anti-rat Alexa Fluor 594 (Invitrogen, A-11007), goat anti-guineapig Alexa Fluor 647 (Abcam, ab150187), goat anti-rabbit Alexa Fluor 647 (Jackson ImmunoResearch, 111-605-144) were used at 1:500 dilution. Zeiss LSM 980 confocal microscopy with a 20X objective lens was used for wide-field IBA1 imaging, a 63X oil objective lens with Z stack (0.2 µm thickness/slice, total 30-40 slices/image) was used for engulfment imaging, and Airyscan 2 with a 63X oil objective lens was used for super-resolution imaging of vGluT1 and PSD95 colocalization. Zeiss LSM 900 confocal microscopy with a 63X oil objective lens was used for TLR4/IBA1 co-staining imaging. In total, 49 images at 3 hours and 52 images at 24 hours post-injection were analyzed for IBA1 fluorescence intensity, 20 images were analyzed for IBA1/TLR4 colocalization, 47 images were analyzed for microglia engulfment, and 26 images at 24 hours and 61 images at 2 weeks post-injection were analyzed for vGluT1/PSD95 colocalization. Images were randomly selected from cortex regions of brain slices. Immunostaining and imaging were performed under same conditions and same settings at same time across different groups within same cohort. Cohort 1 with 8 groups was for IBA1 staining. Cohort 2 with 2 groups for TLR4/IBA1 co-staining. Cohort 3 with 2 groups was for IBA1/CD68/vGluT1 co-staining. Cohort 4 with 2 groups was for 24 hours post-injection vGluT1/PSD95 co-staining. Cohort 5 with 4 groups was for 2 weeks post-injection vGluT1/PSD95 co-staining. Biological triplicates of brain slices from one mouse brain were employed. Results from at least three mouse brains per group were analyzed and compared. ImageJ software was used for IBA1 fluorescence intensity quantification. Imaris software (Oxford Instrument) was used for microglia engulfment 3D rendering analysis. ZEN Blue software was used for colocalization analysis. See details on 3D rendering and colocalization analysis protocols in the Supplementary Information.

### Western blot

Western blot was performed as previously described ^94^. Whole cell lysates from mouse forebrain hemisphere were prepared using 1x lysis buffer (Cell Signaling Technology, 9803S) containing 1x protease/phosphatase inhibitor (Cell Signaling Technology, 5872S). Crude postsynaptic density (PSD) fraction from mouse forebrain hemisphere were prepared following previously described protocol with some modifications^35,95^. See details in Supplementary Information. Samples were boiled in 4x Laemlli buffer (Bio-Rad, 161-0747) at 98 °C for 5 min before gel electrophoresis. Home-made anti-rabbit SHANK3 C-terminus monoclonal antibody (1:1000), anti-rabbit TLR4 monoclonal antibody (1:1000, Santa Cruz, sc-293072), anti-mouse PSD95 monoclonal antibody (1:1000, Invitrogen, MA1-046), HOMER 1b/c monoclonal HRP-conjugated antibody (1:1000, Santa Cruz, sc-17842-HRP), anti-rabbit GluA1 (AMPA receptor) polyclonal antibody (1:1000, Abcam, ab31232), anti-NR2A (NMDA receptor) polyclonal antibody (1:1000, Millipore, ab31232), and β-ACTIN monoclonal HRP-conjugated antibody (1:1000, Invitrogen, MA5-15739-HRP) were used.

### Statistical analysis

Graphpad Prism 10.0 (GraphPad Software) was used for the statistical analysis and data plot. Differences were analyzed by student’s *t*-test if comparing two variables or using ANOVA if comparing more than two variables followed by multiple comparisons either using statistical hypothesis testing with correction or planned comparisons without correction. Significance was determined by two-tailed *t*-test *p* value or multiplicity adjusted *p* value. For data, which was not sampled from Gaussian distribution, a nonparametric test was performed. Since unequal variance observed in LPS-evoked cytokines and chemokines, Brown-Forsythe and Welch ANOVA tests were performed (**Fig. 4A-B**). See detailed statistical analysis for each figure in **Supplementary Table 5**. n represents the total number of data points per group, usually meaning the mice number with an exception that n represents number of images in immunohistochemistry results. Both sexes of mice were used and balanced across genotype and treatment, with an exception that WT+PBS and WT+LPS groups were all male mice in RNA-seq experiments. Behavioral difference between male and female mice were explored in **Supplementary Fig. 1**. In box plot, individual data points were plotted as: whiskers go down to the smallest value and up to the largest value, box extends from the 25th to 75th percentiles and line in the middle of the box is plotted at the median. In bar graph, individual data points were plotted and presented as average ± SEM. *p < 0.05* was considered statistically significant. BioRender was used to create schematic diagram.

## Authorship contribution statement

S-N.Q. and Y-H.J. designed the experiments. S-N.Q. collected and analyzed the data.

S.W. manually scored grooming behavior. S.W., K-Y.K. and S.J. performed western blot.

S-N.Q. and Y-H.J. wrote and revised the manuscripts.

## Acknowledgement

This work was supported by the National Institutes of Health (NIH) grant HD088007 MH104316, MH098114, MH117289, HD087795 (Y-H.J), the Autism Science Foundation (ASF) fellowship (S-N.Q), and the National Research Foundation of Korea (NRF) grant (RS-2024-00401971 (S.J), Brain Pool program RS-2024-00409006 (S.E.W)). This work was supported by the rodent behavior analysis facility and imaging core facility in Yale University funded by the Kavli Institute of Neuroscience. The bulk RNA sequencing was supported by the National Institute of General Medical Sciences of the National Institutes of Health under Award Number 1S10OD030363-01A1.

