## Supplementary information for "Inflammation increases the penetrance of behavioral impairment in *Shank3* haploinsufficiency mice – can it explain the behavioral regression in Autism?"

**Supplementary methods:**

**Bulk RNA sequencing**

*RNA-Seq Quality Control*: Total RNA quality is determined by estimating the A260/A280 and A260/A230 ratios by nanodrop. RNA integrity is determined by running an Agilent Bioanalyzer or Fragment Analyzer gel, which measures the ratio of the ribosomal peaks. Samples with RIN values of 7 or greater are used for library prep.

*RNA-Seq Library Prep*: mRNA is purified from a normalized input between 50-1000ng of total RNA with oligo-dT beads and sheared by incubation at 94°C in the presence of $Mg^{2+}$ (Kapa mRNA Hyper Prep). Following first-strand synthesis with random primers, second strand synthesis and A-tailing are performed with dUTP for generating strand-specific sequencing libraries. Adapter ligation with 3' dTMP overhangs are ligated to library insert fragments. Library amplification amplifies fragments carrying the appropriate adapter sequences at both ends. Strands marked with dUTP are not amplified. Indexed libraries that meet appropriate cut-offs for both are quantified by RT-qPCR (KAPA Biosystems) and insert size distribution determined with the LabChip GX or Agilent TapeStation. Samples with a yield of ≥0.5 ng/ul are used for sequencing.

*Flow Cell Preparation and Sequencing*: Sample concentrations are normalized to 2.0 nM and loaded onto an Illumina NovaSeq flow cell at a concentration that yields 25 million passing filter clusters per sample. Samples are sequenced using 100bp paired-end sequencing on an Illumina NovaSeq according to Illumina protocols. The 10bp unique dual index is read during additional sequencing reads that automatically follow the completion of read 1. Data generated during sequencing runs are simultaneously transferred to the YCGA high-performance computing cluster. A positive control (prepared bacteriophage Phi X library) provided by Illumina is spiked into every lane at a concentration of 0.3% to monitor sequencing quality in real time.

*Data Analysis and Storage*: Signal intensities are converted to individual base calls during a run using the system's Real Time Analysis (RTA) software. Base calls are transferred from the machine's dedicated personal computer to the Yale High Performance Computing cluster via a 1 Gigabit network mount for downstream analysis. Primary analysis - sample de-multiplexing and alignment to the mouse genome - is performed using Illumina's CASAVA 1.8.2 software suite. The data are returned to the user if the sample error rate is less than 2% and the distribution of reads per sample in a lane is within reasonable tolerance.

**Preparation of crude PSD-1 fraction**

Crude PSD-1 fraction was prepared followed previous protocol[1]. Protease and phosphatase inhibitors were used throughout. Forebrain hemisphere tissues were homogenized in HEPES-buffered sucrose (0.32 M sucrose, 4 mM HEPES, pH 7.4) and centrifuged at 800 × g for 10 min at 4°C. The cloudy supernatants were collected and transferred to a new set of tubes. After 12,000 × g centrifugation for 15 min at 4°C, the pellet (P2) and the supernatant were separated. The P2 fraction was lysed using water, then buffered with HEPES (pH 7.4) to 4 mM, and the sample was mixed by rotation at 4°C for 30 min, followed by centrifugation at 20,500 × g for 30 min to yield the synaptosomal membrane (SPM) fraction. The SPM was resuspended in a buffer containing 50 mM HEPES (pH 7.4), 2 mM EDTA, and 0.5% Triton X- 100. After 15 min of mixing by rotation at 4°C, the crude PSD-1 fraction was obtained by centrifugation at

21,130 × g for 20 min at 4°C. The crude PSD pellet was dissolved in 1% SDS-PBS for further quantitative immunoblot analysis.

**Immunohistochemistry analysis**

Microglia engulfment 3D rendering followed previous protocol[2,3]. Briefly, z-stack of high-resolution confocal images of co-staining of IBA1, CD68 and vGlut1 were collected using a 63x oil objective len with 2x zoom to capture single microglia. Then Imaris 10.2 software was used for following process. Median filter was applied to IBA1 channel. Microglia in IBA1 channel was surface rendered with 0.1 µm smoothing. Then background subtraction at 10 µm local contrast, threshold intensity at 1000 and then filtering out surface with volume smaller than 5 $\mu m^3$. A masked CD68 channel was generated using IBA1 surface to isolate the CD68 signals only within microglia. Masked CD68 channel was surface rendered with 0.01 µm smoothing. Then background subtraction at 5 µm local contrast, threshold intensity at 1000 and then filtering out surface with volume smaller than 0.02 $\mu m^3$. A masked vGluT1 channel was generated using CD68 surface to isolate the vGluT1 signals only within CD68 ($vGluT1_{IBA1+CD68+)}$. A second masked vGluT1 channel was generated using IBA1 surface to isolate the vGluT1 signals only within IBA1 ($vGluT1_{IBA1+}$). Masked vGluT1 channel was surface rendered with 0.005 µm smoothing. Then background subtraction at 5 µm local contrast, threshold intensity at 4000 and then filtering out surface with volume smaller than 0.005 $\mu m^3$. Surface material for IBA1 and CD68 was transparent one and for vGluT1 was texture non-transparent one. The total volume of reconstructed surface of IBA1, CD68 and vGluT1 was collected. CD68 occupancy ($IBA1^+$) was calculated as: volume of CD68/volume of IBA1. vGluT1 occupancy ($IBA1^+$) was calculated as: volume of $vGluT1_{IBA1+}$/volume of IBA1. vGluT1 occupancy ($IBA1^+CD68^+$) was calculated as: volume of $vGluT1_{IBA1+CD68+}$/volume of CD68.

Colocalization was performed on a pixel-by-pixel basis in ZEN blue 3.12 software. Every pixel in the image was plotted in the scatter diagram based on its intensity level from each channel. With setting the threshold in each channel, a scatter plot with four quadrants was generated. Same threshold was used across all images in one cohort. Pixels in colocalized quadrant were used for generating white “Colocalization mask” images and further analysis. Software automatically measured colocalization coefficient (Pearson’s coefficient), normalized coefficient, and fluorescence intensity in each channel. Colocalization coefficient normalized by individual channel was calculated as sum of intensity of all pixels in colocalized quadrant divided by sum of intensity of all pixels above the threshold value in specific channel ($\frac{\sum SumGreyCh_{1colocalized}}{\sum SumGreyCh_{1total}}$). Fluorescence intensity was calculated as sum of all gray values from channel divided by the total number of pixels in this channel ($\frac{\sum GreyCh1_i}{AreaCh1}$).

**Supplementary Figures:**

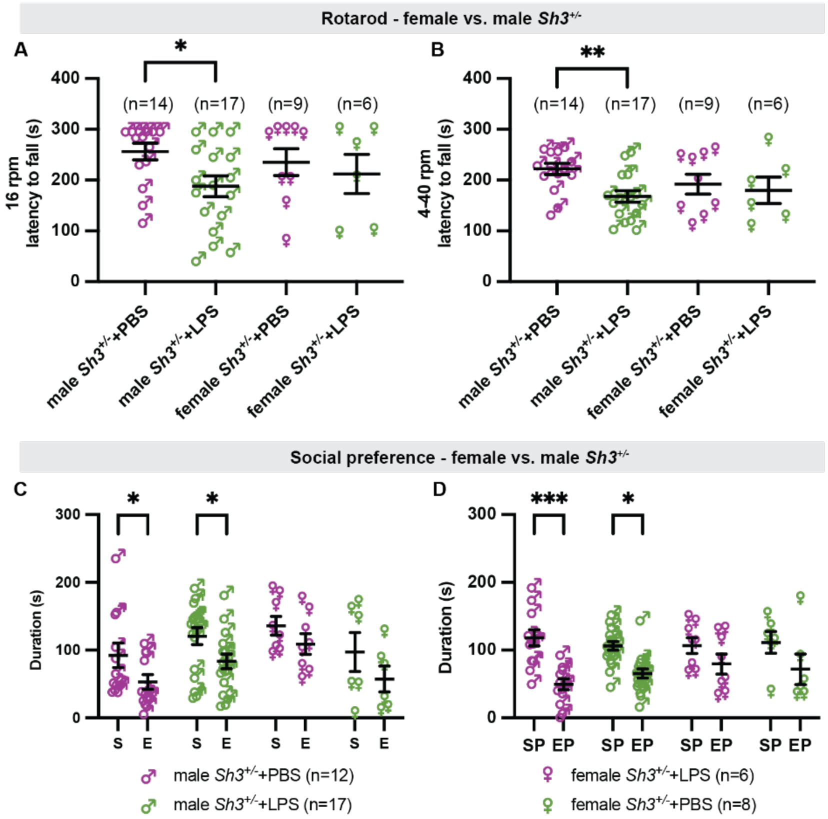


**Supplementary Figure 1. Comparison of LPS effect on motor function and sociability of male and female *Sh3*$^{+/-}$ mice. A-B.** Male *Sh3*$^{+/-}$ +LPS group showed significant reduced latency to fall compared PBS group (16 rpm, $p$ = 0.0197, 4-40 rpm, $p$ = 0.0042). **C-D**. Duration spent in mouse chamber (S), empty chamber (E), proximity regions in mouse chamber (SP) and empty chamber (EP) in three-chamber tests were compared between PBS and LPS group in both female and male *Sh3*$^{+/-}$ mice**.** Two-way repeated measures ANOVA was performed (**C.** sex effect $F_{(3, 39)}$ = 3.367, $p$ = 0.0281, S/E effect $F_{(1, 39)}$ = 11.67, $p$ = 0.0015, sex X S/E interaction $F_{(3, 39)}$ = 0.0768, $p$ = 0.9721; **D.** sex effect $F_{(3, 39)}$ = 0.8197, $p$ = 0.4909, SP/EP effect $F_{(1, 39)}$ = 18.24, $p$ = 0.0001, sex X SP/EP interaction $F_{(3, 39)}$ = 0.8294, $p$ = 0.4858). Both LPS and PBS male *Sh3*$^{+/-}$ mice spent significant longer duration in S vs. E (PBS, $p = 0.038$, LPS, $p = 0.0207$) or SP vs. EP (PBS, $p = 0.0005$, LPS, $p = 0.0103$). $^{*}$ $p < 0.05$, $^{**}$ $p < 0.005$, $^{***}$ $p < 0.0005$.

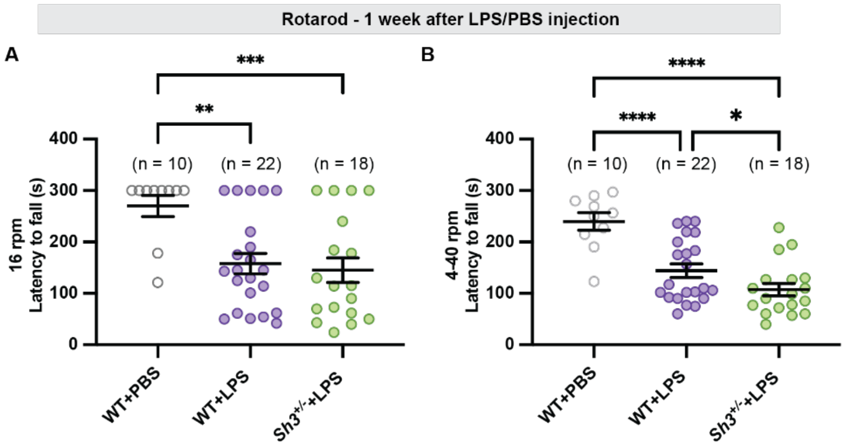


**Supplementary Figure 2. Both WT and *Sh3*$^{+/-}$ mice showed impaired motor function at 1-week post LPS injection compared to WT+PBS group. A-B.** Compared to WT+PBS group, decreased latency to fall was observed in WT+LPS (16 rpm: $p = 0.0024$, 4-40 rpm: $p < 0.0001$) and *Sh3*$^{+/-}$ +LPS (16 rpm: $p = 0.0012$, 4-40 rpm: $p < 0.0001$). *Sh3*$^{+/-}$ +LPS showed slightly more severe motor impairment in 4-40 rpm rotarod test than WT+LPS group ($p = 0.045$). $^{*}$ $p < 0.05$, $^{**}$ $p < 0.005$, $^{***}$ $p < 0.0005$, $^{****}$ $p < 0.0001$.

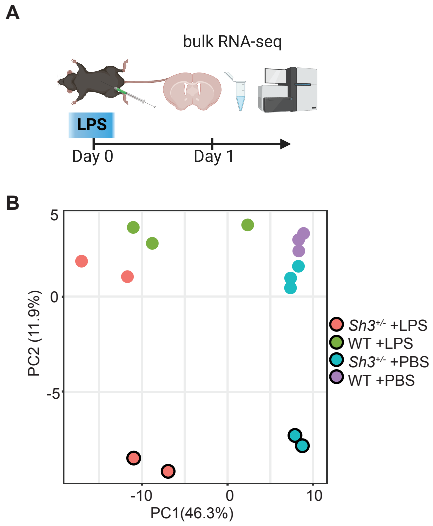


**Supplementary Figure 3. Schematic drawing for RNA-seq experiment design and Principal Component Analysis (PCA) plot of the sequencing results.** Four groups of mice forebrains were sequenced: WT+PBS (n = 3, 3 male mice), WT+LPS (n = 3, 3 male mice), *Sh3*$^{+/-}$+PBS (n = 6, 3 male and 3 female mice), *Sh3*$^{+/-}$+LPS (n = 4, 2 male and 2 female mice). Female mice samples are indicated by circles with black border and male mice samples are indicated by circles without border.

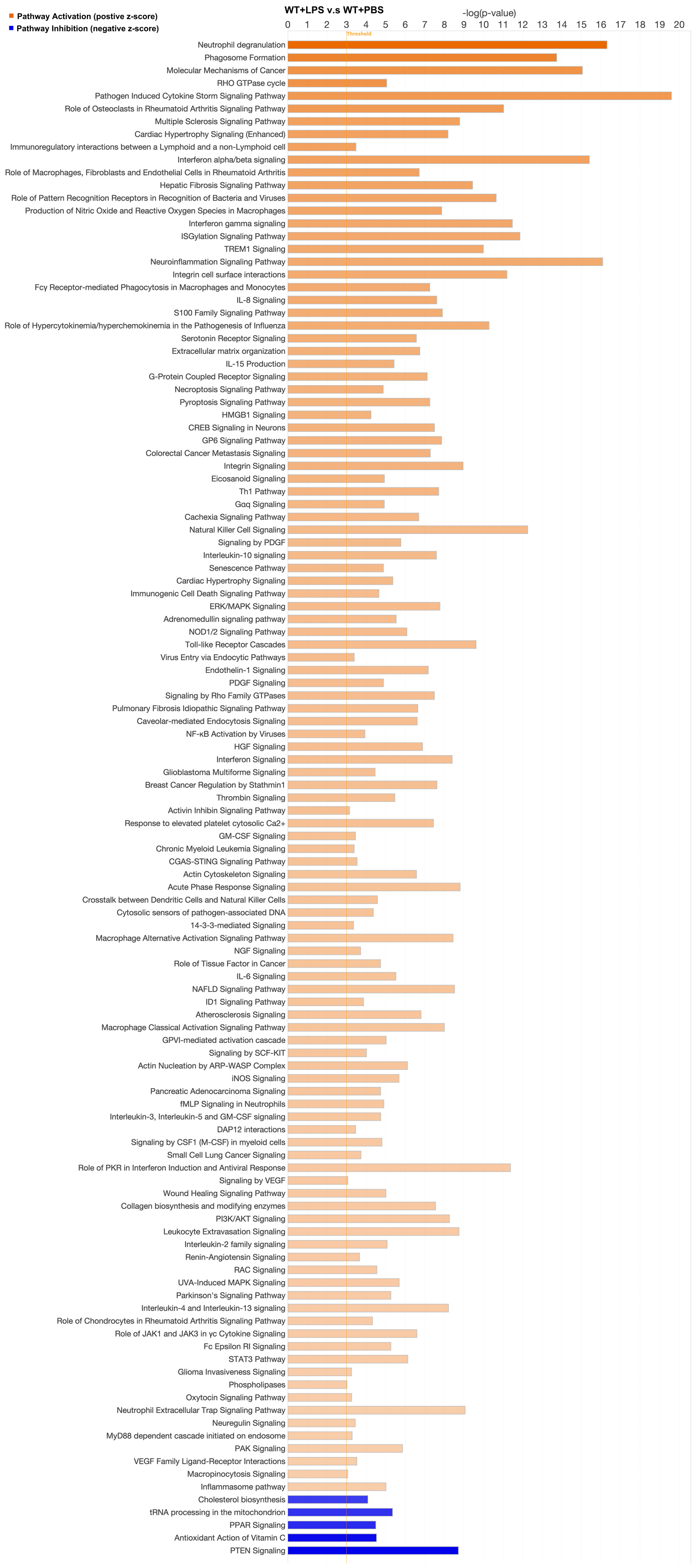
Pathway Activation (postive z-score)
Pathway Inhibition (negative z-score)
WT+LPS v.s WT+PBS
-log(p-value)
0 1 2 3 4 5 6 7 8 9 10 11 12 13 14 15 16 17 18 19 20
Threshold
Neutrophil degranulation
Phagosome Formation
Molecular Mechanisms of Cancer
RHO GTPase cycle
Pathogen Induced Cytokine Storm Signaling Pathway
Role of Osteoclasts in Rheumatoid Arthritis Signaling Pathway
Multiple Sclerosis Signaling Pathway
Cardiac Hypertrophy Signaling (Enhanced)
Immunoregulatory interactions between a Lymphoid and a non-Lymphoid cell
Interferon alpha/beta signaling
Role of Macrophages, Fibroblasts and Endothelial Cells in Rheumatoid Arthritis
Hepatic Fibrosis Signaling Pathway
Role of Pattern Recognition Receptors in Recognition of Bacteria and Viruses
Production of Nitric Oxide and Reactive Oxygen Species in Macrophages
Interferon gamma signaling
ISGylation Signaling Pathway
TREM1 Signaling
Neuroinflammation Signaling Pathway
Integrin cell surface interactions
Fcγ Receptor-mediated Phagocytosis in Macrophages and Monocytes
IL-8 Signaling
S100 Family Signaling Pathway
Role of Hypercytokinemia/hyperchemokinemia in the Pathogenesis of Influenza
Serotonin Receptor Signaling
Extracellular matrix organization
IL-15 Production
G-Protein Coupled Receptor Signaling
Necroptosis Signaling Pathway
Pyroptosis Signaling Pathway
HMGB1 Signaling
CREB Signaling in Neurons
GP6 Signaling Pathway
Colorectal Cancer Metastasis Signaling
Integrin Signaling
Eicosanoid Signaling
Th1 Pathway
Gαq Signaling
Cachexia Signaling Pathway
Natural Killer Cell Signaling
Signaling by PDGF
Interleukin-10 signaling
Senescence Pathway
Cardiac Hypertrophy Signaling
Immunogenic Cell Death Signaling Pathway
ERK/MAPK Signaling
Adrenomedullin signaling pathway
NOD1/2 Signaling Pathway
Toll-like Receptor Cascades
Virus Entry via Endocytic Pathways
Endothelin-1 Signaling
PDGF Signaling
Signaling by Rho Family GTPases
Pulmonary Fibrosis Idiopathic Signaling Pathway
Caveolar-mediated Endocytosis Signaling
NF-κB Activation by Viruses
HGF Signaling
Interferon Signaling
Glioblastoma Multiforme Signaling
Breast Cancer Regulation by Stathmin1
Thrombin Signaling
Activin Inhibin Signaling Pathway
Response to elevated platelet cytosolic Ca2+
GM-CSF Signaling
Chronic Myeloid Leukemia Signaling
CGAS-STING Signaling Pathway
Actin Cytoskeleton Signaling
Acute Phase Response Signaling
Crosstalk between Dendritic Cells and Natural Killer Cells
Cytosolic sensors of pathogen-associated DNA
14-3-3-mediated Signaling
Macrophage Alternative Activation Signaling Pathway
NGF Signaling
Role of Tissue Factor in Cancer
IL-6 Signaling
NAFLD Signaling Pathway
ID1 Signaling Pathway
Atherosclerosis Signaling
Macrophage Classical Activation Signaling Pathway
GPVI-mediated activation cascade
Signaling by SCF-KIT
Actin Nucleation by ARP-WASP Complex
iNOS Signaling
Pancreatic Adenocarcinoma Signaling
fMLP Signaling in Neutrophils
Interleukin-3, Interleukin-5 and GM-CSF signaling
DAP12 interactions
Signaling by CSF1 (M-CSF) in myeloid cells
Small Cell Lung Cancer Signaling
Role of PKR in Interferon Induction and Antiviral Response
Signaling by VEGF
Wound Healing Signaling Pathway
Collagen biosynthesis and modifying enzymes
PI3K/AKT Signaling
Leukocyte Extravasation Signaling
Interleukin-2 family signaling
Renin-Angiotensin Signaling
RAC Signaling
UVA-Induced MAPK Signaling
Parkinson's Signaling Pathway
Interleukin-4 and Interleukin-13 signaling
Role of Chondrocytes in Rheumatoid Arthritis Signaling Pathway
Role of JAK1 and JAK3 in γc Cytokine Signaling
Fc Epsilon RI Signaling
STAT3 Pathway
Glioma Invasiveness Signaling
Phospholipases
Oxytocin Signaling Pathway
Neutrophil Extracellular Trap Signaling Pathway
Neuregulin Signaling
MyD88 dependent cascade initiated on endosome
PAK Signaling
VEGF Family Ligand-Receptor Interactions
Macropinocytosis Signaling
Inflammasome pathway
Cholesterol biosynthesis
tRNA processing in the mitochondrion
PPAR Signaling
Antioxidant Action of Vitamin C
PTEN Signaling

**Supplementary Figure 4. Horizontal bar chart displayed significant canonical pathways identified from WT+LPS vs. WT+PBS comparison.** Bar plot of the negative log of p-value (X-axis) vs. the pathway names (Y-axis). p-value is calculated using the right-tailed Fisher's Exact Test. A negative log of p-value cutoff of 1.3 ($p < 0.05$) is used to identify significantly changed functional pathways. The orange bars indicate z-score above 0.1, predicting pathway activation and blue bars indicate z-score below -0.1, indicate predicted inhibition. The bars are sorted so that pathways with highest z-score are on the top of the chart and pathways with lowest z-score are on the bottom of the chart. Pathways with no activity prediction or ineligible for analysis are excluded.

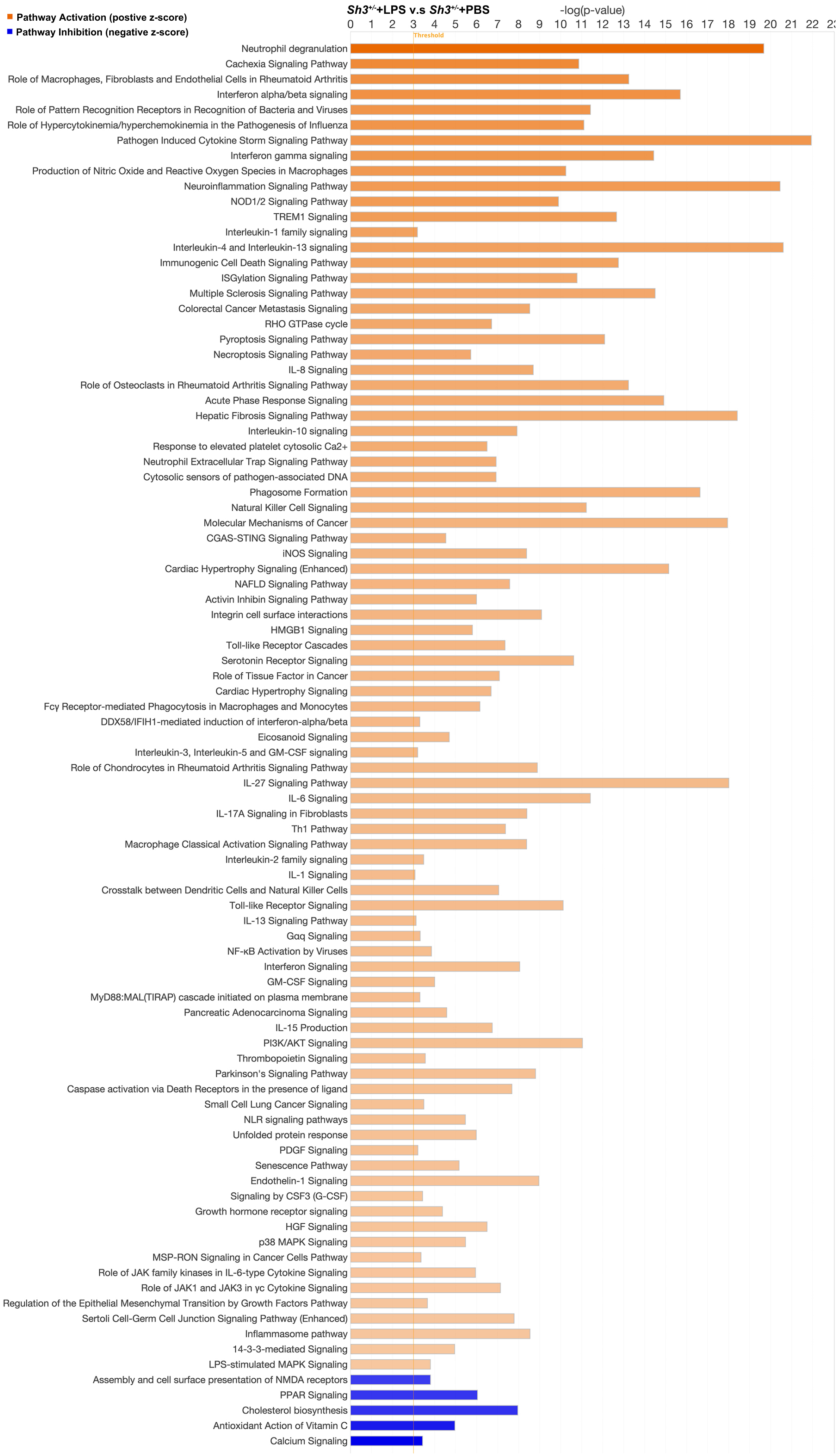
Pathway Activation (postive z-score)
Pathway Inhibition (negative z-score)
Sh3+/-+LPS v.s Sh3+/-+PBS
-log(p-value)
0 1 2 3 4 5 6 7 8 9 10 11 12 13 14 15 16 17 18 19 20 21 22 23
Threshold
Neutrophil degranulation
Cachexia Signaling Pathway
Role of Macrophages, Fibroblasts and Endothelial Cells in Rheumatoid Arthritis
Interferon alpha/beta signaling
Role of Pattern Recognition Receptors in Recognition of Bacteria and Viruses
Role of Hypercytokinemia/hyperchemokinemia in the Pathogenesis of Influenza
Pathogen Induced Cytokine Storm Signaling Pathway
Interferon gamma signaling
Production of Nitric Oxide and Reactive Oxygen Species in Macrophages
Neuroinflammation Signaling Pathway
NOD1/2 Signaling Pathway
TREM1 Signaling
Interleukin-1 family signaling
Interleukin-4 and Interleukin-13 signaling
Immunogenic Cell Death Signaling Pathway
ISGylation Signaling Pathway
Multiple Sclerosis Signaling Pathway
Colorectal Cancer Metastasis Signaling
RHO GTPase cycle
Pyroptosis Signaling Pathway
Necroptosis Signaling Pathway
IL-8 Signaling
Role of Osteoclasts in Rheumatoid Arthritis Signaling Pathway
Acute Phase Response Signaling
Hepatic Fibrosis Signaling Pathway
Interleukin-10 signaling
Response to elevated platelet cytosolic Ca2+
Neutrophil Extracellular Trap Signaling Pathway
Cytosolic sensors of pathogen-associated DNA
Phagosome Formation
Natural Killer Cell Signaling
Molecular Mechanisms of Cancer
CGAS-STING Signaling Pathway
iNOS Signaling
Cardiac Hypertrophy Signaling (Enhanced)
NAFLD Signaling Pathway
Activin Inhibin Signaling Pathway
Integrin cell surface interactions
HMGB1 Signaling
Toll-like Receptor Cascades
Serotonin Receptor Signaling
Role of Tissue Factor in Cancer
Cardiac Hypertrophy Signaling
Fcγ Receptor-mediated Phagocytosis in Macrophages and Monocytes
DDX58/IFIH1-mediated induction of interferon-alpha/beta
Eicosanoid Signaling
Interleukin-3, Interleukin-5 and GM-CSF signaling
Role of Chondrocytes in Rheumatoid Arthritis Signaling Pathway
IL-27 Signaling Pathway
IL-6 Signaling
IL-17A Signaling in Fibroblasts
Th1 Pathway
Macrophage Classical Activation Signaling Pathway
Interleukin-2 family signaling
IL-1 Signaling
Crosstalk between Dendritic Cells and Natural Killer Cells
Toll-like Receptor Signaling
IL-13 Signaling Pathway
Gαq Signaling
NF-κB Activation by Viruses
Interferon Signaling
GM-CSF Signaling
MyD88:MAL(TIRAP) cascade initiated on plasma membrane
Pancreatic Adenocarcinoma Signaling
IL-15 Production
PI3K/AKT Signaling
Thrombopoietin Signaling
Parkinson's Signaling Pathway
Caspase activation via Death Receptors in the presence of ligand
Small Cell Lung Cancer Signaling
NLR signaling pathways
Unfolded protein response
PDGF Signaling
Senescence Pathway
Endothelin-1 Signaling
Signaling by CSF3 (G-CSF)
Growth hormone receptor signaling
HGF Signaling
p38 MAPK Signaling
MSP-RON Signaling in Cancer Cells Pathway
Role of JAK family kinases in IL-6-type Cytokine Signaling
Role of JAK1 and JAK3 in γc Cytokine Signaling
Regulation of the Epithelial Mesenchymal Transition by Growth Factors Pathway
Sertoli Cell-Germ Cell Junction Signaling Pathway (Enhanced)
Inflammasome pathway
14-3-3-mediated Signaling
LPS-stimulated MAPK Signaling
Assembly and cell surface presentation of NMDA receptors
PPAR Signaling
Cholesterol biosynthesis
Antioxidant Action of Vitamin C
Calcium Signaling

**Supplementary Figure 5. Horizontal bar chart displayed significant canonical pathways identified from *Sh3*$^{+/-}$+LPS vs. *Sh3*$^{+/-}$+PBS comparison.**

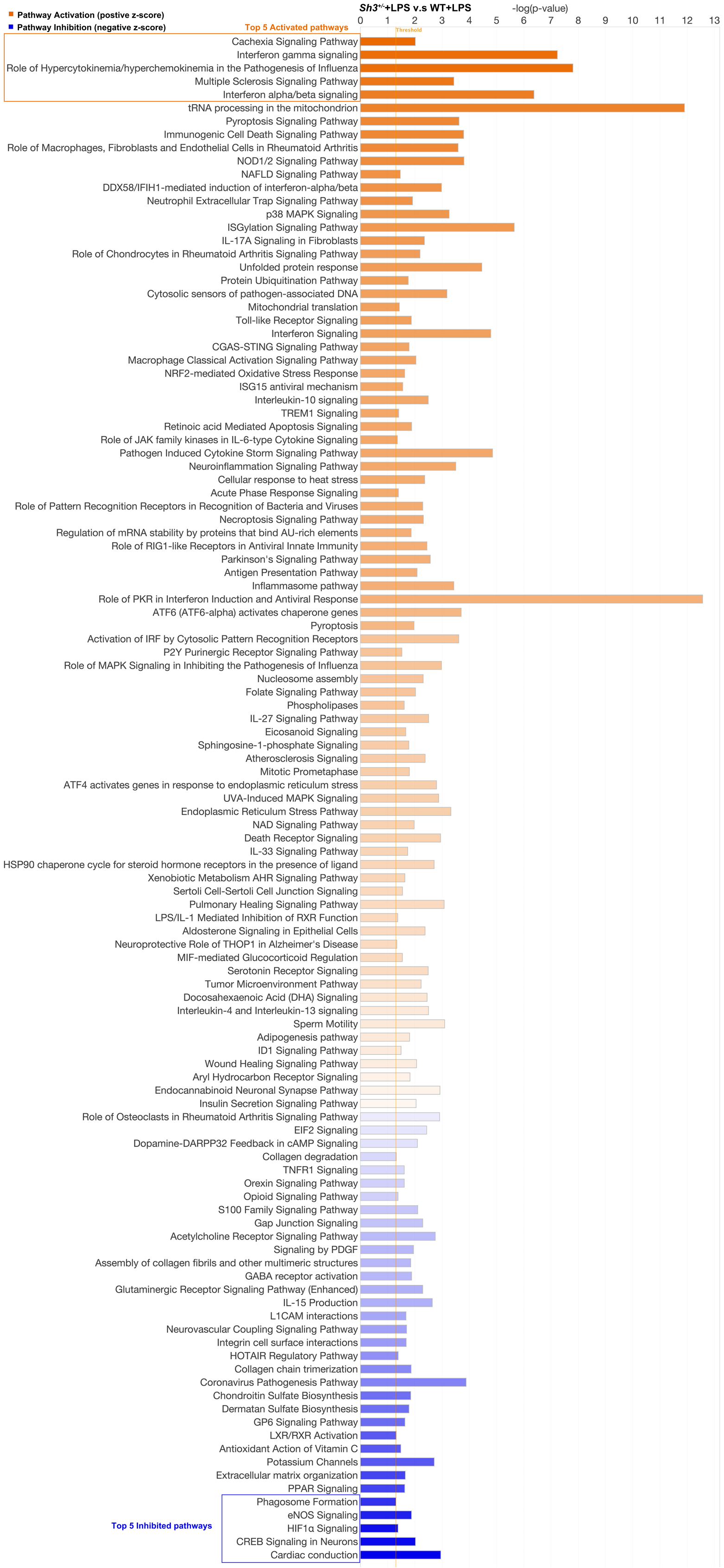

Pathway Activation (postive z-score)
Pathway Inhibition (negative z-score)
Sh3+/-+LPS v.s WT+LPS
-log(p-value)
0 1 2 3 4 5 6 7 8 9 10 11 12 13
Threshold
Top 5 Activated pathways
Cachexia Signaling Pathway
Interferon gamma signaling
Role of Hypercytokinemia/hyperchemokinemia in the Pathogenesis of Influenza
Multiple Sclerosis Signaling Pathway
Interferon alpha/beta signaling
tRNA processing in the mitochondrion
Pyroptosis Signaling Pathway
Immunogenic Cell Death Signaling Pathway
Role of Macrophages, Fibroblasts and Endothelial Cells in Rheumatoid Arthritis
NOD1/2 Signaling Pathway
NAFLD Signaling Pathway
DDX58/IFIH1-mediated induction of interferon-alpha/beta
Neutrophil Extracellular Trap Signaling Pathway
p38 MAPK Signaling
ISGylation Signaling Pathway
IL-17A Signaling in Fibroblasts
Role of Chondrocytes in Rheumatoid Arthritis Signaling Pathway
Unfolded protein response
Protein Ubiquitination Pathway
Cytosolic sensors of pathogen-associated DNA
Mitochondrial translation
Toll-like Receptor Signaling
Interferon Signaling
CGAS-STING Signaling Pathway
Macrophage Classical Activation Signaling Pathway
NRF2-mediated Oxidative Stress Response
ISG15 antiviral mechanism
Interleukin-10 signaling
TREM1 Signaling
Retinoic acid Mediated Apoptosis Signaling
Role of JAK family kinases in IL-6-type Cytokine Signaling
Pathogen Induced Cytokine Storm Signaling Pathway
Neuroinflammation Signaling Pathway
Cellular response to heat stress
Acute Phase Response Signaling
Role of Pattern Recognition Receptors in Recognition of Bacteria and Viruses
Necroptosis Signaling Pathway
Regulation of mRNA stability by proteins that bind AU-rich elements
Role of RIG1-like Receptors in Antiviral Innate Immunity
Parkinson's Signaling Pathway
Antigen Presentation Pathway
Inflammasome pathway
Role of PKR in Interferon Induction and Antiviral Response
ATF6 (ATF6-alpha) activates chaperone genes
Pyroptosis
Activation of IRF by Cytosolic Pattern Recognition Receptors
P2Y Purinergic Receptor Signaling Pathway
Role of MAPK Signaling in Inhibiting the Pathogenesis of Influenza
Nucleosome assembly
Folate Signaling Pathway
Phospholipases
IL-27 Signaling Pathway
Eicosanoid Signaling
Sphingosine-1-phosphate Signaling
Atherosclerosis Signaling
Mitotic Prometaphase
ATF4 activates genes in response to endoplasmic reticulum stress
UVA-Induced MAPK Signaling
Endoplasmic Reticulum Stress Pathway
NAD Signaling Pathway
Death Receptor Signaling
IL-33 Signaling Pathway
HSP90 chaperone cycle for steroid hormone receptors in the presence of ligand
Xenobiotic Metabolism AHR Signaling Pathway
Sertoli Cell-Sertoli Cell Junction Signaling
Pulmonary Healing Signaling Pathway
LPS/IL-1 Mediated Inhibition of RXR Function
Aldosterone Signaling in Epithelial Cells
Neuroprotective Role of THOP1 in Alzheimer's Disease
MIF-mediated Glucocorticoid Regulation
Serotonin Receptor Signaling
Tumor Microenvironment Pathway
Docosahexaenoic Acid (DHA) Signaling
Interleukin-4 and Interleukin-13 signaling
Sperm Motility
Adipogenesis pathway
ID1 Signaling Pathway
Wound Healing Signaling Pathway
Aryl Hydrocarbon Receptor Signaling
Endocannabinoid Neuronal Synapse Pathway
Insulin Secretion Signaling Pathway
Role of Osteoclasts in Rheumatoid Arthritis Signaling Pathway
EIF2 Signaling
Dopamine-DARPP32 Feedback in cAMP Signaling
Collagen degradation
TNFR1 Signaling
Orexin Signaling Pathway
Opioid Signaling Pathway
S100 Family Signaling Pathway
Gap Junction Signaling
Acetylcholine Receptor Signaling Pathway
Signaling by PDGF
Assembly of collagen fibrils and other multimeric structures
GABA receptor activation
Glutaminergic Receptor Signaling Pathway (Enhanced)
IL-15 Production
L1CAM interactions
Neurovascular Coupling Signaling Pathway
Integrin cell surface interactions
HOTAIR Regulatory Pathway
Collagen chain trimerization
Coronavirus Pathogenesis Pathway
Chondroitin Sulfate Biosynthesis
Dermatan Sulfate Biosynthesis
GP6 Signaling Pathway
LXR/RXR Activation
Antioxidant Action of Vitamin C
Potassium Channels
Extracellular matrix organization
PPAR Signaling
Top 5 Inhibited pathways
Phagosome Formation
eNOS Signaling
HIF1α Signaling
CREB Signaling in Neurons
Cardiac conduction

**Supplementary Figure 6. Horizontal bar chart displayed significant canonical pathways identified from *Sh3*$^{+/-}$+LPS vs. WT+LPS comparison.** Top five activated canonical pathways with highest z-score and top five inhibited canonical pathways with lowest z-score are highlighted.

**Supplementary Tables:**

**Supplementary Table 1. Upregulated genes list (see attached excel file)**

**Supplementary Table 2. Downregulated genes list (see attached excel file)**

**Supplementary Table 3. Shared upregulated genes list (see attached excel file)**

**Supplementary Table 4. Shared downregulated genes list (see attached excel file)**

**Supplementary Table 5. Complete statistics (see attached excel file)**
